# Antibody maturation increases rigidity in protein-contacting regions and flexibility at glycan interfaces

**DOI:** 10.64898/2026.05.27.728205

**Authors:** Shahlo O. Solieva, Joyce Park, Justin Miller, Austin Kriews, Amelia Escolano, Gregory R. Bowman

## Abstract

Antibody design is a challenging task that could be improved by understanding the conformational changes accompanying affinity maturation. Antibody maturation is a critical immune system process by which antibodies gain mutations that improve affinity and specificity for antigen targets. However, how these mutations change antibody dynamics, specifically whether antibodies become more rigid as they mature, remains a contested topic. Using adaptive sampling molecular dynamics simulations and over 8.5 milliseconds of Folding@home simulations of seven lineages, we discover that affinity maturation selectively tunes the dynamics of antibody paratope regions depending on whether they contact protein residues or glycans. We find that antibody regions that contact glycans evolve to become more flexible and, consistent with other studies, antibody regions that contact protein residues on the antigen become rigid. This pattern holds regardless of whether one includes the constant region in the simulations, indicating that the computational cost of all-atom antibody MD simulations can be reduced by half without sacrificing the accuracy of the variable region’s dynamics. We expect the principles identified in this study will enable precise, dynamics-based engineering of high-affinity antibodies and to inform immunogen design against challenging, glycan-shielded viral targets such as HIV.

**Significance Statement:** Antibodies are currently one of the most powerful therapeutics available. Our bodies evolve antibodies to bind a target (called an antigen) through a process called maturation. Understanding how antibody dynamics change during maturation would lead to improved antibody design strategies. Through extensive all-atom molecular dynamics simulations of multiple antibody lineages, we demonstrate that the dynamics accompanying affinity maturation are dependent on the antigen binding site. Antibody regions that contact protein residues on the antigen become rigid while regions that contact glycans gain flexibility with maturation. These findings can be leveraged for improved, dynamics-based antibody design.

## Introduction

Antibodies are powerful therapeutics due to their ability to bind antigens with high specificity and affinity, yet antibody design remains a major challenge. The high conformational heterogeneity and sequence diversity of the complementarity-determining region (CDR) loops enable broad antigen recognition but complicate structure prediction^1^. While advances in structure determination have led to extensive collections of antibody structures^2^, these structures represent only static snapshots of the conformational landscapes explored in solution^3^. Recent AI-based tools such as AlphaFold3 have improved antibody structure prediction but still struggle with CDR loops due to their structural variability and sequence diversity^4,5^. There are also large language models (LLMs) specifically developed for antibody structure prediction, which provide quick predictions but typically only generate one structure per sequence query^6^. While some tools do generate an ensemble of structures, they do not encapsulate the entirety of the conformations explored or provide information on how long antibodies spend at these conformations^5,7^. There are also AI-based tools that have used molecular dynamics (MD) simulation data for fine-tuning, taking protein dynamics into account for improved performance^8^. Taken together, the limitations of these tools highlight the importance of understanding protein conformational landscapes for improved rational protein design.

Understanding how antibody dynamics change during the natural affinity maturation process could provide valuable insights into the criteria necessary for designing high affinity antibodies. The immune system is designed to have high sequence diversity, with humans having the ability to create up to an estimated 10^18^ unique naïve antibody sequences^9^. One major source of this sequence diversity is from gene segments recombining to produce a variable heavy and light chain^10^. Both variable chains have three CDR loops, which are the loops that usually interact the most with antigens^11^. The third CDR in the heavy chain (CDRH3) has the highest sequence and structural diversity. This sequence diversity allows antigen-naïve antibodies to have some level of binding to antigens encountered. Antibodies then achieve high specificity and affinity through a process called affinity maturation, an essential part of the adaptive immune response where through rounds of exposure to an antigen, antibodies gain random mutations that increase their specificity and affinity to the antigen. Modern sequencing advances enable us to track when and where sequence changes occur in antibody lineages during the maturation process, and innovations in structure determination give us valuable insight into how the affinity maturation process can affect antibody structures^12^. While these advances show tremendous progress towards understanding affinity maturation, they do not provide insight into the conformational heterogeneity and dynamic nature of antibodies and proteins in general. Therefore, a significant question remains unclear – how do the mutations accompanying affinity maturation affect antibody dynamics and can these dynamics be exploited to guide in silico antibody design?

The current, conventional paradigm of affinity maturation is that antibodies become rigid during this process to optimize contacts and increase shape complementarity to the target antigen. One of the earliest demonstrations of this rigidification hypothesis is from an X-ray crystallography study in 1997, where Wedemeyer et al. found that the hapten-binding affinity-matured antibody 48G7 had a smaller amount of conformational change when bound to hapten compared to its antigen-naïve antibody^13^. In the years since, there have been numerous studies suggesting a rigidification of the antigen-interacting regions or CDR loops in antibodies with affinity maturation^14–23^. These studies encapsulate a diverse set of methods including X-ray crystallography, cryogenic electron microscopy (cryo-EM), surface plasmon resonance, three-pulse photon echo peak shift spectroscopy, isothermal calorimetry, and hydrogen-deuterium exchange mass spectrometry.

While the dominant explanation of affinity maturation has been the rigidification theory, there is a more recent and growing body of evidence that suggests affinity maturation is much more complex. For instance, a comprehensive study of the antibody structural database by Jeliazkov et al. did not find differences in the flexibility of the CDRH3 loops from naïve and mature antibodies^24^. They instead conclude that rigidification of the CDRH3 loop is only one of the mechanisms selected for and therefore is not a consistent result of affinity maturation. Other studies suggest a variety of mechanisms that could be responsible for affinity maturation, such as CDR loops or other regions gaining flexibility and shifts in the orientation between the variable heavy and light chain domains^25–28^. These studies show the current uncertainty surrounding the role of antibody dynamics in affinity maturation.

Molecular dynamics (MD) simulations offer powerful, atomic-level insights into protein dynamics. Likely due to the high computational cost of running MD simulations, there are not many studies that have investigated antibody maturation using large sets of all-atom MD simulations. Consistent with the experimental results outlined previously, existing MD studies have reached multiple conclusions. A number of MD simulation studies of antibody lineages have observed that affinity maturation results in rigidification of CDR loops^29–32^. A recent large-scale study of coarse-grained antibody simulations found naïve antibodies were more flexible on average than mature antibodies, but this difference was not statistically significant and a clear rigidification trend with maturation was not found^33^. In contrast to this rigidification paradigm other studies suggest an increase in dynamics with affinity maturation, two examples from MD simulation studies being the broadly neutralizing anti-HIV antibody 3BNC60 and the anti-GD2 antibody 3F8^25,26^. Additionally, there is a study which found that the heavy chain in three antibody lineages becomes rigid while the light chain gains flexibility^34^. These differing conclusions could represent a real diversity in mechanisms for achieving high affinity. They could also represent statistical artifacts, as some studies use as little as 10 ns of simulation but much longer timescale processes are likely relevant. Much of the published work has also made simplifying assumptions for the sake of computational expediency. For example, it is common to leave out the constant regions to minimize the system size. It would be valuable to assess how reasonable this is by comparing to simulations that include the constant regions. Many studies also propose trends based on comparing the naïve and fully mature antibodies without considering any of the intermediates that arose during the maturation process.

Here, we present extensive simulations of seven antibody lineages, including multiple intermediates when possible, to assess whether there are trends in the rigidity of different regions during maturation. In total, we ran over 8.5 milliseconds of aggregate simulation data, which is roughly two orders of magnitude more data than the most extensive all atom MD studies we are aware of^29^. The constant domains were included in the simulations, making up the fragment antigen-binding (Fab) antibody. For a subset of antibodies, we simulated the isolated variable region (fragment variable, Fv) and compared the results with simulations that include the constant region.

## Results

### There is no correlation between changes in global flexibility and maturation

To explore the relationship between protein dynamics and maturation, we chose to simulate 31 antibody sequences from seven different lineages (see Tables S1 and S2 for sequences). The antibody lineages we include in this study are anti-HIV antibodies CH103, CAP256, 1-18, PGT121, EPTC112^35–38^, anti-influenza antibody CH65^39,40^, and anti-GD2 antibody 3F8^41^. These lineages were selected on the basis of having one or more of the following: multiple known intermediate sequences, published structures of the antibodies bound to their respective antigens, published structures of their naïve antibodies, published binding affinity values, or due to them being ideal candidates that others have also studied. We chose to include multiple intermediates to track any trends that may develop during affinity maturation. While the naïve and mature antibodies from each lineage provide information about the antibody at the start and end of affinity maturation, the intermediate antibodies may help bridge the gap between these endpoints. Additionally, many of the past MD studies only simulated the variable region of antibodies, raising questions about the constant region’s effect on the variable region. We included the constant region in our simulations. We used FAST^42^ simulations to study antibody dynamics throughout the affinity maturation process. FAST, fluctuation amplification of specific traits, is a goal-oriented sampling method that allows us to increase our chances of seeing fluctuations along a collective variable of interest without introducing perturbations to the potential energy function. It has been shown to outperform conventional simulations by at least an order of magnitude^42^. The collective variable we used in our simulations is a root-mean-squared deviation (RMSD) maximization of the CDRH3 loop from its starting structure. We chose the CDRH3 loop due to its high structural diversity^43^. While the other five loops (CDRL1-3 and CDRH1-2) have been shown to adopt canonical conformations^44,45^, the CDRH3 is very difficult to classify structurally, therefore, we reasoned that RMSD maximization of this loop should be prioritized to drive the conformational exploration of the antibodies being simulated. Once each antibody was simulated, we clustered the data based on the RMSD of the variable heavy and light chains, then created Markov state models (MSM) (see Methods). We were then left with representative cluster centers that we used for further analysis and visualization. The representative cluster centers and associated MSM probabilities for all antibodies discussed in this manuscript are available on the OSF database. We calculated per-residue alpha carbon root-mean-squared fluctuation (RMSF) values for each antibody by finding the Euclidean distance between the XYZ coordinates of each representative cluster center to the average XYZ position of all representative cluster centers for that antibody, with cluster centers weighted by their MSM equilibrium probabilities (see Methods). These per-residue alpha carbon RMSF values will be referred to simply as ‘RMSF’ going forward. We then ran simulations on Folding@home^46^ to corroborate our FAST simulations for the naïve and mature antibodies from five lineages and find that the results are similar despite having an almost 100-fold difference in aggregate simulation time (Figure S1). We collected a total of over 8.5 milliseconds of simulation data from Folding@home and over 100 microseconds from FAST.

Our simulations show that some lineages gain rigidity during maturation while others gain flexibility. Variable region RMSF data do not show a collective, general trend in global flexibility with affinity maturation across the seven lineages. There are lineages that show a decrease in global variable region RMSF with maturation (CH65, CH103, CAP256) and lineages that show an increase (EPTC112, 3F8, 1-18, PGT121), although not all of these differences are statistically significant according to the 95% bootstrapped confidence intervals (Figure 1A) (see Methods for confidence interval calculations). The most populated states from each MSM encompassing 20% of the conformations visited from simulations of CH103 and PGT121 are shown in Figure 1B to illustrate the decrease and increase, respectively, in variable region RMSF with maturation. As mentioned previously, in their study of three antibody lineages, Li et al. observed that while there was not a global trend in dynamics with affinity maturation, the variable heavy chains did become rigid with maturation, while the light chains became flexible^34^. To determine if this trend was also present in our lineages, we calculated the average RMSF values of the variable heavy and light chains separately. However, we still do not see a trend across all lineages when comparing only variable heavy or light chains (Figure S2). We then wanted to see whether intermediate antibodies in these lineages have RMSF values that fall between the naïve and mature values. We find that the intermediates do not necessarily follow the trend set by the naïve and mature endpoint antibodies (Figure S2). There are lineages where some of the intermediates do not follow the trend; this is the most apparent in the 1-18 lineage, where intermediate I1 has RMSF values that are significantly higher (no overlap in 95% confidence intervals) than the corresponding naïve and mature antibodies (Figure 1C, D). The sequence for the 1-18 intermediate I1 was created through a combinatorial analysis of antibodies belonging to the same family from the same donor^37^ and is closest in sequence identity to naïve 1-18 out of the three intermediates included in this study. We find that the global and per-residue RMSF values from the FAST and Folding@home simulations generally agree (Figure S1), indicating that FAST can reliably capture the conformational landscapes of the antibodies in this study. Therefore, we are confident that the FAST simulations accurately capture the dynamics of the intermediate antibodies and other lineages investigated.

**Figure 1.**
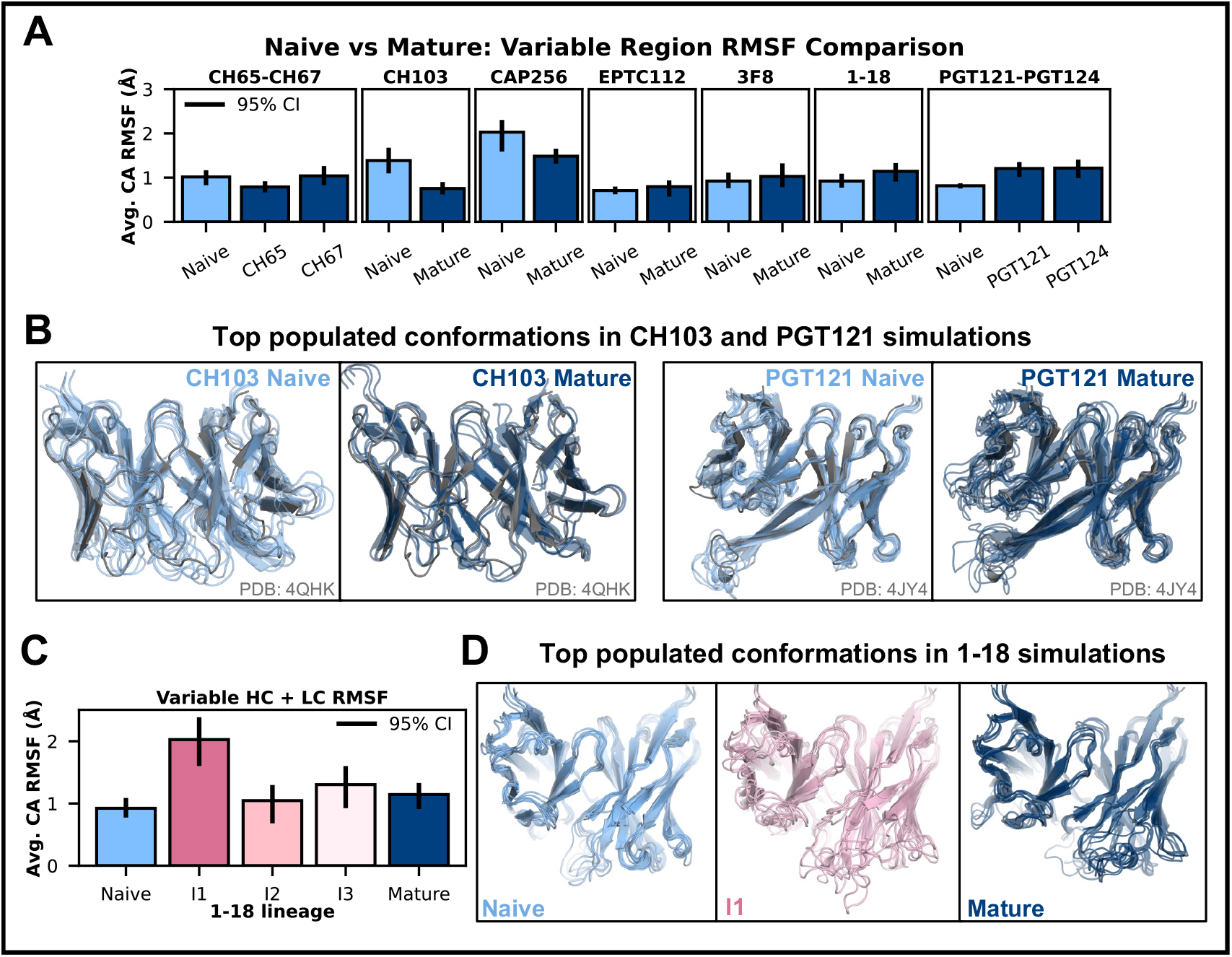
The global RMSF does not have a clear trend with maturation. **(A)** For the variable chain (including both the heavy chain and light chain), alpha carbon RMSF values for each pair of naïve (light blue) and mature (dark blue) antibodies do not show a consistent trend across all lineages. The error bars represent a 95% confidence interval, see Methods for how these values were calculated. **(B)** Top 20% most populated conformations from simulations of a lineage that becomes rigid (CH103) with maturation and a lineage that becomes flexible (PGT121) with maturation, focused on the variable heavy and light chains. For both examples, the light blue structures represent the naïve antibody, dark blue structures represent the mature antibody, and gray structures represent a crystal structure from the corresponding antibody lineage (PDB 4QHK and 4JY4) **(C)** Considering three intermediates from the 1-18 lineage makes it clear that there is no monotonic trend with maturation. The error bars represent a 95% confidence interval. The three intermediates used are described in Table S1. **(D)** The top 20% most populated conformations from 1-18 naïve (light blue), I1 intermediate (pink), and mature (dark blue) simulations show the higher degree of structural fluctuations in the I1 intermediate antibody compared to the naïve and mature antibodies.

### Results are robust to removing the constant region or using an alternative light chain

Whether the constant region influences the dynamics of the variable region remains an unanswered question with direct implications on antibody engineering. As mentioned previously, many MD studies reduced their computational cost by only simulating the variable regions of antibodies. However, it remains unclear if this is reasonable or if including the constant region would dramatically alter the results. To determine whether the absence of the constant region affects the dynamics of the variable region, we ran FAST simulations of the variable regions for three antibody lineages, including 1 to 5 intermediates per lineage, and compared them to our simulations that include the constant region.

We find that the behavior of the variable region is largely independent of whether the constant region is included. The global and per-residue RMSF values show that the presence of the constant region does not significantly affect the dynamics of the variable region in 13 of the 14 simulations (Figure 2A). The one exception is the I10 intermediate of the CAP256 lineage, where the absence of the constant region resulted in a significantly higher RMSF. These results suggest there is not substantial allostery between the variable and constant regions and that conclusions based on isolated variable regions will generally apply to the full-length antibody.

**Figure 2.**
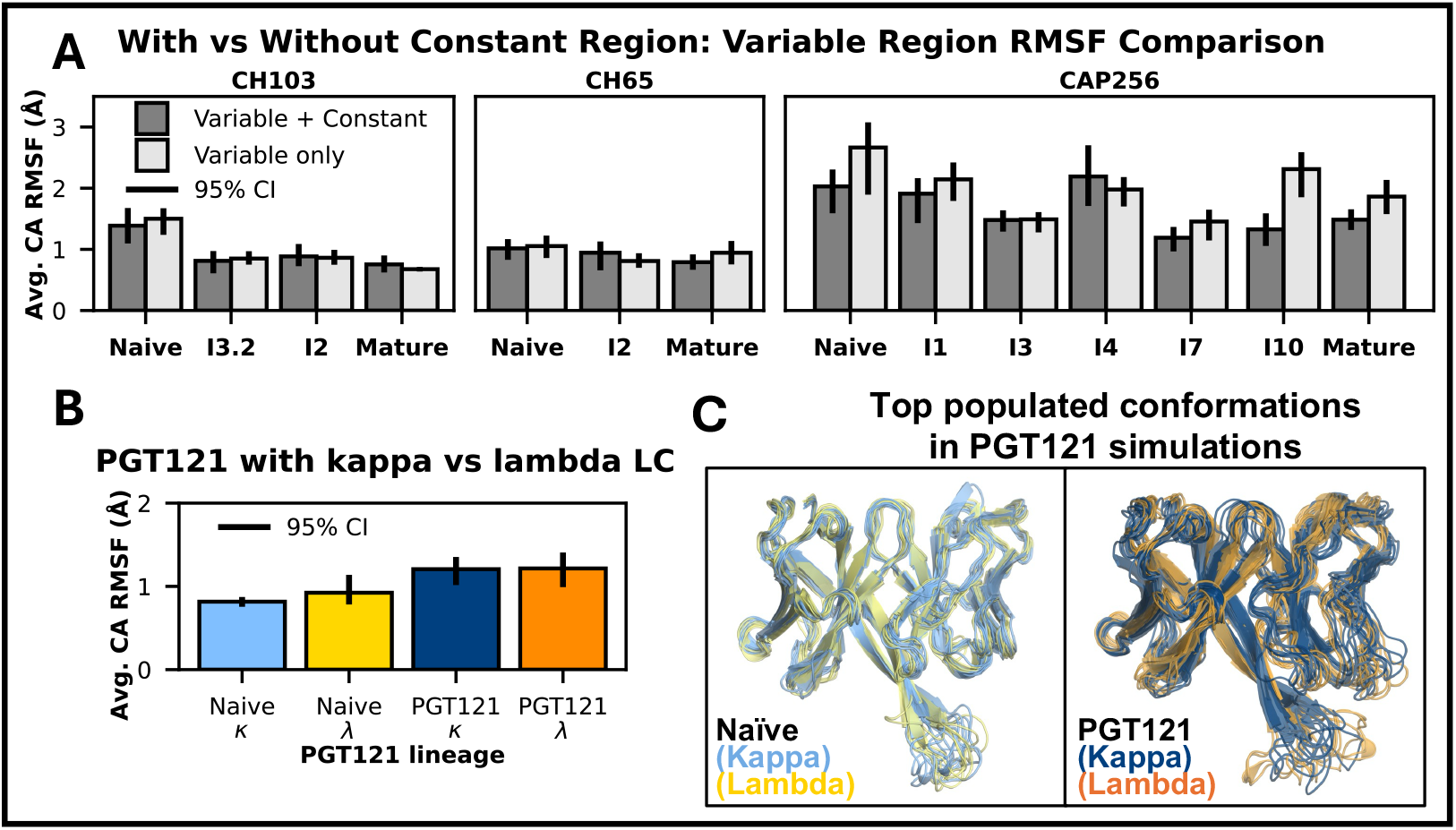
The variable region behaves similarly in the presence and absence of the constant region and in the case of PGT121, with alternative light chain constants. **(A)** The average variable region (both heavy and light chains) alpha carbon RMSF values are not significantly different between simulations that include the constant region (dark gray) and those without (light gray), as indicated by the overlapping 95% confidence intervals represented by the error bars, for all but one set of simulations (CAP256 I10). **(B)** The average variable heavy chain and light chain RMSF values for PGT121 naïve and mature with Kappa vs with Lambda light chains do not show significant differences, indicated by the overlapping 95% confidence intervals represented by the error bars. PGT121 naïve and mature antibodies with the Kappa light chain constant are shown in light blue and dark blue, respectively. PGT121 naïve and mature antibodies with the Lambda light chain constant are shown in yellow and orange, respectively. **(C)** The top 20% most populated conformations from simulations of PGT121 naïve antibody with Kappa (light blue) vs Lambda (yellow) constant chains and PGT121 mature antibody with Kappa (dark blue) vs Lambda (orange) constant chains show that the degree of structural fluctuations remains similar when the light chain constants are swapped.

We explored the effects of the constant region further by swapping the Kappa and Lambda light chain constants in an antibody lineage, anticipating that this change in the constant region would not affect the variable region dynamics. The constant region in the light chain can be encoded by two different chromosomes, resulting in either a Kappa or Lambda light chain isotype, with the Kappa light chain naturally being more prevalent in healthy humans^47^. We ran simulations of the naïve and mature antibodies of the PGT121 lineage with either the Lambda or Kappa light chain constant.

Upon global and per-residue RMSF analysis, we found no significant difference in their global variable region dynamics and in most residues (Figure 2B, 2C, S3). Similarly, no major effects on production or antigen binding were found in a study where the light chain constant region isotype was swapped in two anti-Her2 therapeutic antibodies^48^.

Taken together, these results suggest that only simulating the variable region, and thus reducing computational cost by at least half, is sufficient to study the dynamics of the variable region without sacrificing accuracy. However, information about the dynamics of the constant region is still relevant for antibody engineering, as the flexibility of the constant region and the elbow angle between the variable and constant regions are shown to be influenced by the identity of the light chain^49^. Additionally, reversion of mutations in the elbow region of a mature antibody has been shown to decrease its potency by altering the conformational flexibility of the interdomain and paratope regions^50^. From an engineering perspective, it would be valuable to test whether manipulating the angle between the variable and constant regions by changing light chain isotype or introducing elbow region mutations could result in higher affinity antibodies from either reducing unfavorable or promoting favorable antigen interactions.

### Antibody regions that contact antigen residues become rigid with maturation

The lack of a consistent trend in global antibody dynamics with affinity maturation does not eliminate the possibility of there being trends within localized regions across the lineages. As described previously, several studies reported a preorganization of loop conformations at antibody paratopes with maturation to aid antigen binding through a ‘lock-and-key’ mechanism^29^. Therefore, we reasoned that loops contacting the antigen, such as the CDRH3 in many antibodies, may also display this preorganization or rigidification in our simulations.

We focused on the antigen-contacting regions at antibody paratopes to investigate how their dynamics change with affinity maturation. For each antibody lineage, we used a published structure of the mature antibody bound to its antigen to define antibody residues interacting with protein residues on the antigen (see Methods for the PDB ID codes). We calculated the minimum distance of every antibody heavy atom to every antigen heavy atom in the mature crystal or cryo-EM structures. We used a cutoff of 6 Å between heavy atoms to define an antibody residue as interacting with a protein residue on the antigen. Figure S4 shows per-residue RMSF profiles with protein-contacting residues marked with gray points. Neighboring residues were grouped into ‘regions’ for simplicity and are shown in Figures S5-11 for lineages CH103, CH65, CAP256, 1-18, PGT121, 3F8, and EPTC112 with plots zoomed in on each region. In some cases, these regions overlap with CDR loops.

The simulations show that affinity maturation broadly results in rigidification of antibody residues that contact only protein residues on the antigen. This trend is most evident in lineages CH103 and CH65, which have eight and four protein-only contacting regions with a significant decrease in RMSF with maturation (Figure S5 and S6). Significance here is determined by whether there is overlap in the 95% confidence intervals of the naïve and mature antibody RMSF values. If there is no overlap in at least one of the protein-contacting residues in a given region, then the difference in that region is considered significant. Figure 3 illustrates the RMSF decrease in lineage CH65’s naïve and mature variable heavy chains. The three protein-only contacting regions all fall within the CDRH loops (Figure 3B, Table S2). The crystal structure of mature CH65 bound to its antigen, influenza hemagglutinin, with a focus on the three protein-only contacting regions is shown in Figure 3C. The most populated MSM states that encompass 20% of the conformational states visited from simulations of naïve and mature CH65, aligned to the mature crystal structure, show the conformational heterogeneity in the naïve antibody and the rigidification of these regions in the mature antibody (Figure 3D). Interestingly, four of the total twelve mutations in the heavy chain occur directly at the antigen-contacting residues and show a significant decrease in RMSF. Two of these mutations are of glycine, a very flexible amino acid, to aspartate, which explains the rigidity gained. The aspartate may have been selected to improve electrostatic interactions with the antigen. The other two mutations are of tyrosine and asparagine to histidine, which may again have been selected to improve electrostatic interactions. Lineage 1-18 has two protein-only contacting regions, where one region, HC Region 4, shows a significant decrease with maturation, while the other region, HC Region 2, has a mix of residues showing significant decreases and increases in flexibility (Figure S8A). Taken together, these results, in line with previous studies, suggest that antibodies evolve to reduce conformational entropy at residues that interact only with protein residues on the antigen, likely to maximize or stabilize favorable, binding-competent contacts with the antigen.

**Figure 3.**
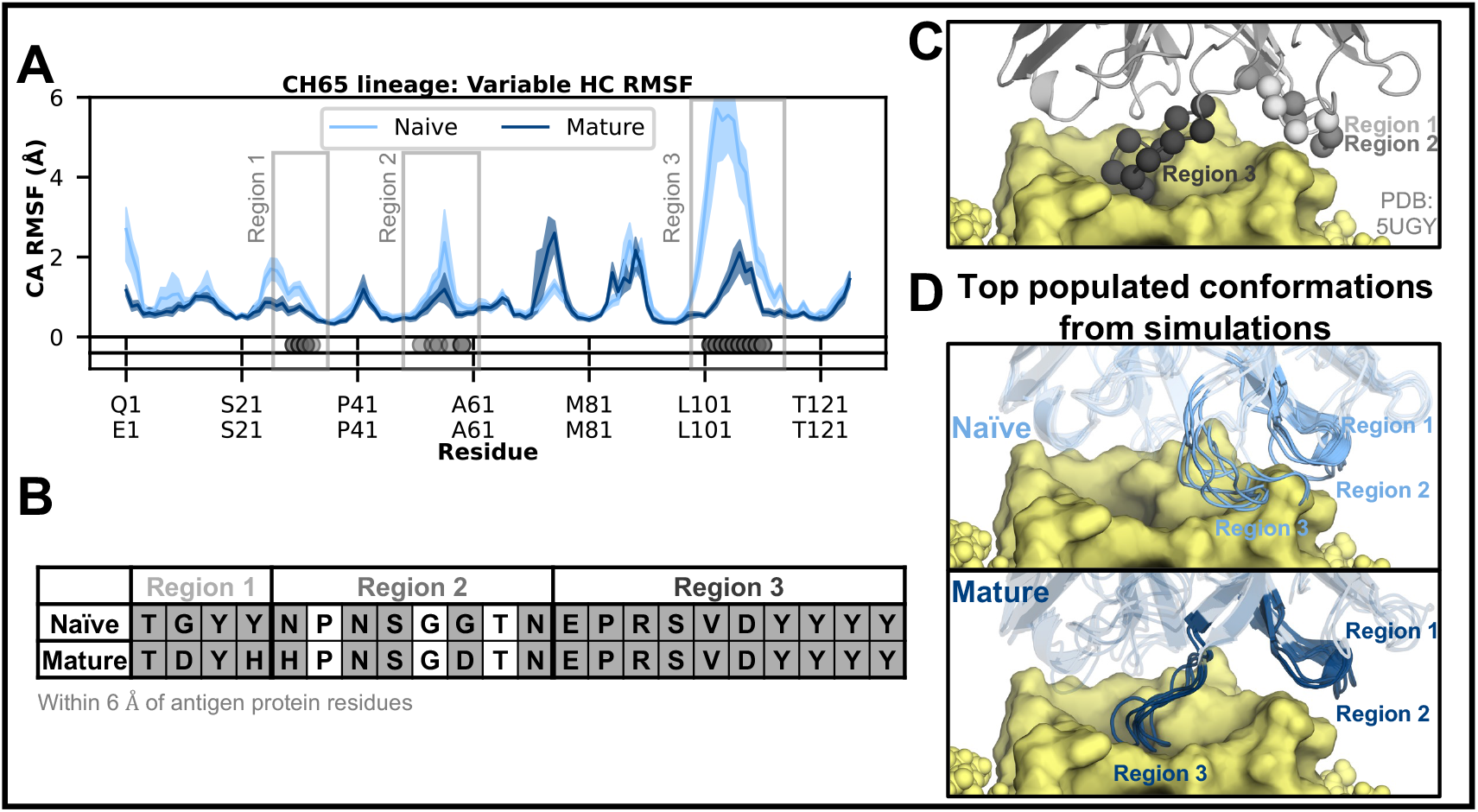
Antibody residues that interact with protein residues on the antigen become more rigid during maturation. The CH65 lineage is used for illustration. **(A)** Sequence-aligned per-residue RMSF plot of the heavy chain of the naïve (light blue) and mature (dark blue) antibodies, with gray dots in the first row under the plot at antibody residues within 6 Å of the protein antigen residues in the mature PDB structure (PDB: 5UGY). The dots are transparent if the RMSF difference between the naïve and mature simulation at that residue is not significant (overlap in the confidence intervals). These residues are grouped into three regions, as outlined by the gray boxes. The second row under the plot the plot is reserved for marking residues interacting with glycans, but there are none in this case. The shaded outlines represent a 95% confidence interval. The x-axis ticks are labeled with the aligned residue sequence for the naïve (top row) and mature (bottom row) antibodies. **(B)** Sequence alignment of the naïve and mature antibodies for the three approximate regions that interact with the antigen, with gray shading at the residues within 6 Å of the antigen protein residues. The three regions all fall within the CDRH1-3 loops. See Table S2 for the full antibody sequences with the CDR loops highlighted. **(C)** The PDB structure of CH65 mature antibody in complex with its antigen (yellow), with antibody residues within 6 Å of the antigen protein residues shown as gray spheres, colored by region: light gray for Region 1, gray for Region 2, dark gray for Region 3. **(D)** The top 20% most populated conformations from simulations of the naïve (transparent light blue) and mature antibody (transparent dark blue), with the three protein antigen interacting regions labeled and colored (light blue for naïve, dark blue for mature).

In contrast, regions that contact both protein residues and antigen glycans do not consistently follow this rigidification trend, suggesting that there are other forces in play. Of the fourteen total regions that contact both, four regions (belonging to lineages CH103 and EPTC112) show a significant decrease in RMSF with maturation, one region (belonging to lineage CAP256) has residues with both significant decreases and increases, seven regions (belonging to lineages 1-18 and PGT121) show a significant increase, and two regions (belonging to lineages EPTC112 and 1-18) do not have any residues with a significant difference.

### Antibody regions that contact glycans become flexible with maturation

Motivated by the complexity of dynamics in the protein and glycan contacting antibody regions, we sought to characterize how glycans influence the dynamics of their binding partners during affinity maturation. The approach we took here was the same as the protein-contacting antibody residue analysis, but with an added focus on the glycan-contacting residues. We repeated the heavy atom to heavy atom distance calculation, but with glycans rather than the antigen protein residues to define a residue within 6 Å as glycan-contacting. The 3F8 antibody lineage does not have a published structure in complex with its GD2 antigen; the residues highlighted as contacting the GD2 antigen are from an in-silico docking study (Figures S4 and S10)^51^. Although the GD2 antigen is a ganglioside and not a true glycan, it does contain a glycan chain, therefore, the 3F8 residues that contact GD2 are considered glycan-contacting residues in this study.

We find that antibody residues contacting only glycans show an increase in flexibility with maturation. Across all seven lineages studied, there are nine regions that contact only glycans. Four of these regions have a significant RMSF difference between the naïve and mature antibodies, three of which show a significant increase in RMSF with maturation (lineages 1-18, 3F8, and PGT121, Figure S4). The remaining region, HC Region 1 of CAP256, shows a significant decrease in RMSF with maturation, which may result from the significant RMSF decrease in HC Region 2, which contacts HC Region 1 (Figure S7). As noted previously, seven of the fourteen regions contacting both protein residues and glycans show an increase in RMSF with maturation. Figure 4A shows the variable light chain of lineage 1-18, which has one glycan-only and two protein-and-glycan contacting regions, all three regions gain flexibility with maturation. Notably, the glycan-only contacting region (HC Region 2) falls within the DE loop, also known as the CDRL4 loop^52^. This loop is understudied, yet its flexibility may reflect a broader role in glycan binding and could be a valuable target for affinity-improving mutations.

**Figure 4.**
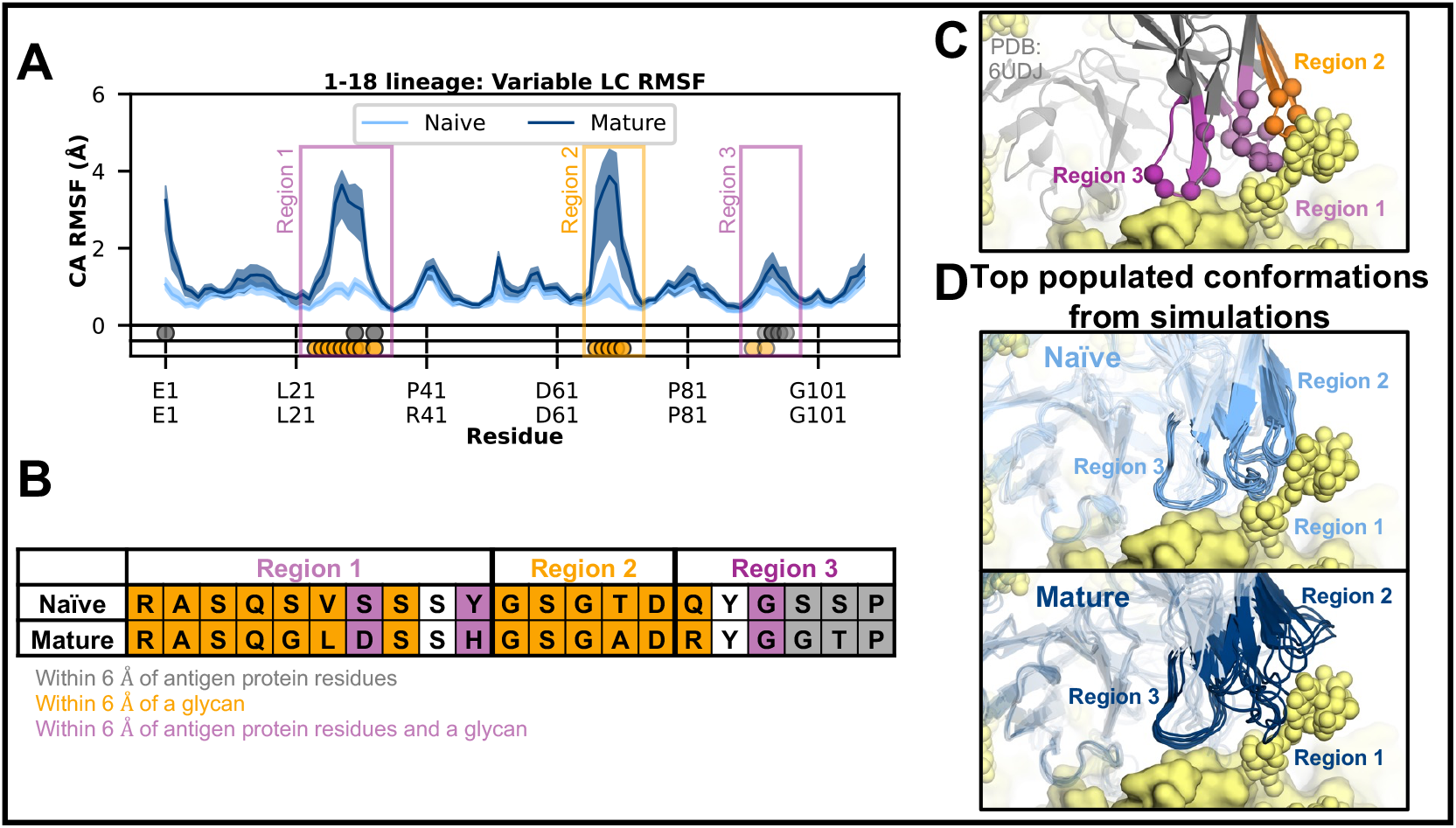
Antibody residues near antigen glycans become more flexible with maturation. The 1-18 lineage is used for illustration. **(A)** Sequence-aligned per-residue RMSF plot of the light chain of the naïve (light blue) and mature (dark blue) antibodies, with shaded regions representing a 95% confidence interval. Gray dots in the first row under the plot are placed at antibody residues within 6 Å of the antigen protein residues in the mature PDB structure (PDB 6UDJ). Similarly, in the second row under the plot, orange dots are placed at antibody residues that are within 6 Å of the antigen glycans. The dots are transparent if the RMSF difference between the naïve and mature simulation at that residue is not significant (overlap in the confidence intervals). The regions that interact with both antigen protein residues and glycans are outlined in purple boxes, the region that interacts only with antigen glycans is outlined in orange. The x-axis ticks are labeled with the aligned residue sequence for the naïve (top row) and mature (bottom row) antibodies. **(B)** Sequence alignment of the naïve and mature antibodies for the three regions outlined in (A), with shading at residues within 6 Å of the antigen protein residues (gray), glycans (orange), or both (purple). The CDRL1 and CDRL3 loops fall within or partially within Region 1 and Region 3, respectfully. Region 2 contains the CDRL4 loop. See Table S2 for the full antibody sequences with the CDR loops highlighted. **(C)** The PDB structure of 1-18 mature antibody in complex with its antigen (PDB 6UDJ), with antibody residues within 6 Å of the antigen protein residues, glycans, or both shown as spheres, colored by region: light purple for Region 1, orange for Region 2, purple for Region 3. The antigen is shown as a yellow surface with glycans as yellow spheres. **(D)** The top 20% most populated conformations from simulations of the naïve (transparent light blue) and mature antibody (transparent dark blue), with the three regions labeled and colored (light blue for naïve, dark blue for mature).

An interesting glycan-only contacting region that undergoes a large, maturation-induced conformational change is the CDRL1 loop of the EPTC112 lineage (Figure 5A). The CDRL1 loop is in an alpha-helical conformation in the HIV Env trimer-bound mature EPTC112 cryo-EM structure (PDB ID: 8C8T) and in the naïve EPTC112 simulations. However, simulations of mature EPTC112 reveal that this helix unfolds, allowing it to extend into solution. Figure 5B shows the top 20% populated conformations from naïve and mature EPTC112, focused on the CDRL1 loop and aligned to the antigen-bound cryo-EM structure of mature EPTC112 (PDB 8C8T). It is possible that the helical unfolding and consequent loop flexibility may have evolved to help facilitate interactions with the nearby N301 glycan to aid binding. The cryo-EM structure shows that the only light chain to antigen contact under a 6 Å distance occurs between the last residue in the CDRL1 loop and the N301 glycan (Figure 5C). However, it is possible that the rest of the CDRL1 loop contacts the N301 glycan, as the full N301 glycan is not resolved in the cryo-EM structure. Therefore, we aligned this structure to one that contains the fully glycosylated HIV Env trimer (PDB ID 5FYL) and found that the rest of the CDRL1 loop can contact the full N301 glycan (Figure 5D)^53^. It is known that the N301 glycan is important for mature EPTC112’s ability to bind to this region, as Molinos-Albert et al. found that removing the N301 glycan abrogates its binding to the HIV Env trimer^38^. Therefore, it is plausible that if EPTC112 evolved to rely on binding to the N301 glycan, one of the mechanisms for binding the N301 glycan is increased flexibility at the CDRL1 loop via the helical unfolding.

**Figure 5.**
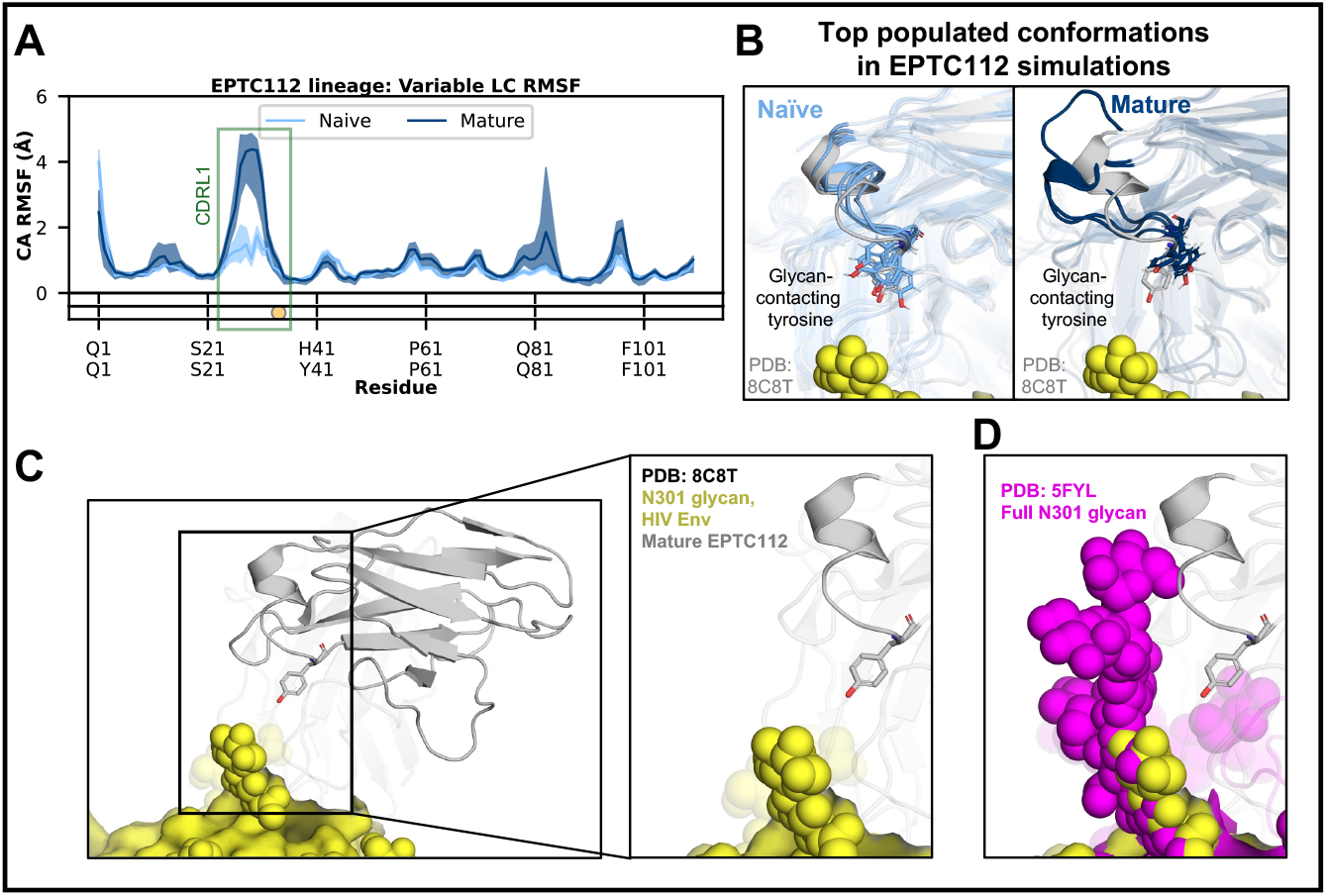
Maturation induces helical unfolding in CDRL1 loop in the EPTC112 lineage. **(A)** Sequence-aligned per-residue RMSF plot of the light chain of the naïve (light blue) and mature (dark blue) antibodies, with shaded regions representing a 95% confidence interval. The orange dot represents the tyrosine residue which is within 6 Å of an antigen glycan in the mature PDB structure (PDB 8C8T), with transparency indicating that the RMSF difference between the naïve and mature simulations is not significant. This glycan-interacting region is outlined with an orange box, while the approximate location of the CDRL1 loop is outlined with a green box. The x-axis ticks are labeled with the aligned residue sequence for the naïve (top row) and mature (bottom row) antibodies. **(B)** The top 20% most populated conformations from simulations of the naïve (transparent light blue) and mature (transparent dark blue) EPTC112 antibodies, aligned to the antigen-bound cryo-EM structure of the mature antibody (gray) in complex with its antigen (yellow) with the glycan depicted as yellow spheres (PDB 8C8T). The CDRL1 loop is highlighted with full opacity, and the glycan-contacting tyrosine residue is shown explicitly. **(C)** Cryo-EM structure of mature EPTC112 (gray) bound to HIV Env trimer (yellow) with the N301 glycan shown as yellow spheres (PDB 8C8T). **(D)** Alignment of PDB 8C8T and 5FYL shows that the full N301 glycan (magenta) can reach the CDRL1 loop in EPTC112.

Together, these results suggest that the affinity maturation process increases flexibility at glycan-contacting regions. Reasons for this increase may be to maintain favorable contacts with highly dynamic glycans and accommodate a range of glycan variants, especially for viral glycoproteins with high glycan heterogeneity, such as gp120 from the HIV envelope^54,55^.

## Discussion

We investigated affinity maturation using all-atom MD simulations of seven antibody lineages. While there was no global trend in flexibility, we did identify consistent flexibility trends in localized regions. The simulation results show that protein-only contacting antibody regions become rigid with maturation while glycan-only contacting regions gain flexibility. These trends suggest that conformational entropy is selectively tuned during affinity maturation at specific paratope regions. Our results also show, consistent with other studies^26^, that intermediates do not always follow the trends set by their naïve and mature counterparts; this is true for global (Figure S4) and local trends (Figures S5B to S10B).

In addition to characterizing the dynamics underlying affinity maturation, this study provides insight into reducing the computational cost of all-atom antibody MD simulations. We demonstrate that, at a much smaller fraction of the cost and time, FAST simulations recapitulate the results (global and local flexibility trends) from 8.5 milliseconds of Folding@home data (Figures S5B to S9B). While generating milliseconds of simulation data is not feasible on a single GPU in a reasonable amount of time, a FAST run can be completed on a GPU in a matter of days, depending on the system size and simulation length. Furthermore, we show that the global and local dynamics of the variable region are generally not affected by the presence of the constant region (Figures S5B to S7B). Thus, it would be reasonable to only simulate the variable regions of antibodies, reducing computational cost by at least half without sacrificing accuracy.

The findings outlined in this study have broad implications for in silico antibody engineering. In silico antibody design is an incredibly challenging task, with dynamics being historically underutilized, making it essential to understand how affinity maturation changes the conformational landscapes of antibodies for improvements in affinity. The local flexibility trends identified here could be leveraged as a general principle for the design of higher-affinity antibodies: rigidify protein-only contacting regions or increase flexibility at glycan-only contacting regions. Incorporating this principle into antibody design pipelines could improve the accuracy of antibody-antigen affinity predictions, guide mutation selection to optimize therapeutic antibodies, and inform the design of immunogens to elicit higher-affinity antibodies.

## Materials and Methods

All representative cluster centers, along with MSM equilibrium probabilities per center, are available on the Open Science Framework (OSF, https://osf.io/wn396/overview?view_only=7193b3578df84294be49204d2b639c0e). All simulation, analysis, and visualization code, along with relevant PDB structures, are available on GitHub (https://github.com/ssolieva/antibody-dynamics.git).

### Molecular Dynamics Simulations

The simulations were run using Gromacs 2023.2^56^ and the Charmm36m force field^57^ at 310K. For each antibody, we either used a published PDB structure and modeled in any missing residues using the Modeller tool in UCSF Chimera X^58^ or used AlphaFold3 to model the antibody^5^. If a PDB structure was used, the PDB ID is included in Table S1, the sequences for all antibodies simulated are in Table S2 with CDR loops highlighted^59^. The antibody structure was then placed in a dodecahedron box with a minimum 1 nm distance between the antibody and the edge of the box. It was solvated using the TIP3P water model and 0.1 M NaCl at a neutral charge. The second step was to equilibrate the structure. Energy minimization was performed using steepest decent minimization until the maximum force of the system reached 1000 kJ/mol/nm or until 50,000 steps were performed. NVT equilibration was performed for a total of 200 ps, followed by 1000 ps of NPT equilibration. We used FAST^42^ to maximize the root-mean-squared deviation of the CDRH3 loop (backbone and beta-carbon atoms). We ran 5 generations with 10 kids, each kid being a 50 ns production run, totaling 2500 ns of FAST simulation time for each antibody. Frames were saved every 20 ps. All files for running the simulations are in the Github repository.

### Folding@home Simulations

We ran simulations on Folding@home^46^ to corroborate our FAST simulations for the naïve and mature antibodies from five lineages (CH103, CH65, CAP256, PGT121, 1-18). Simulations were run in OpenMM 8.2.0 using the Charmm36m force field at 310K with a step size of 2 fs, friction of 1 ps^-1^ and a nonbonded cutoff of 1.2 nm. We collected an aggregate of 8.5 milliseconds of all-atom MD simulation data. The simulations were all seeded from the same starting structures used for the FAST simulations. Each antibody system had around 1000 trajectories, with frames saved at 40 ps intervals.

### Clustering and MSM Approach

Simulation clustering was performed using K-hybrid clustering from the ENSPARA software^60^. K-hybrid clustering uses RMSD clustering and stops creating clusters once the maximum RMSD reaches some distance (0.15 nm variable region backbone + CB atom RMSD in this case). It uses a combination of K-centers and K-medoids. K-centers for initialization and K-medoids for refinement. K-centers minimizes the maximum distance between any point and its cluster center. K-medoid uses actual data points (simulation frames) as cluster centers, while K-centers doesn’t necessarily have to use an actual data point. The FAST data was not subsampled, but the Folding@Home data was subsampled by 50 frames due to its large size. A cluster radius of 0.15 nm RMSD was used for all simulations except for CAP256 where the cluster radius was reduced to 0.215 nm to account for increased flexibility.

Markov state model building was also performed using the ENSPARA software. First, implied timescales were built for each system at 100 different lag times between 0.2 ns and 100 ns. The final MSMs were built using the lag time at which the implied timescales plateau, in this case, 1 ns.

### RMSF Calculations

RMSF was calculated by first pre-aligning all MSM center structures using MDTraj^61^. Next, we found the population-weighted average position for each alpha carbon and computed the Euclidean distance each average alpha carbon position and the corresponding alpha carbon position in the MSM center (for each MSM center). This is a per-residue deviation from the average position for each cluster center. Finally, we weighted each center’s alpha carbon distance by the population-weight to achieve per-residue RMSF values. For region-level calculations, the RMSF across the region was averaged.

Bootstrapping was performed by randomly selecting trajectories from the entire dataset with replacement. Existing assignments were re-used and MSMs were re-constructed for each bootstrapped dataset. Each dataset was bootstrapped a total of 1000 times.

### Minimum antibody to antigen distance calculations

The following PDB ID codes were used for these calculations: 6vtt (CAP256), 8c8t (EPTC112), 5ugy (CH65), 7uoj (PGT121), 9d7o (CH103), and 6udj (1-18). Most of these structure files have been edited to remove other antibodies that may be included, glycans that are very far from the antibody, and potentially parts of the antigen that are far away from the antibody-antigen binding interface to reduce computational cost and avoid unnecessary calculations. These edited structure files are included in the GitHub repository. Distances were calculated between the variable heavy and light chains of the antibody to the antigen’s protein residues and glycans. The atoms used are all heavy (non-hydrogen) atoms. The scripts and notebooks for analysis are included in the GitHub repository.

## Acknowledgments

We are grateful to the citizen scientists who participate in Folding@home for helping to generate simulation data. We would like to thank Dr. Maria Belen Palacio for making the mice from which the 1-18 I2 sequence was derived. This work was funded by the National Institutes of Health through NIGMS R35GM152085 (G.R.B.), R01 AI172627 (A.E.) and by the Gates Foundation INV-036995 (A.E.).

