## Supplementary material for "Antibody maturation increases rigidity in protein-contacting regions and flexibility at glycan interfaces": SI Figures and Tables

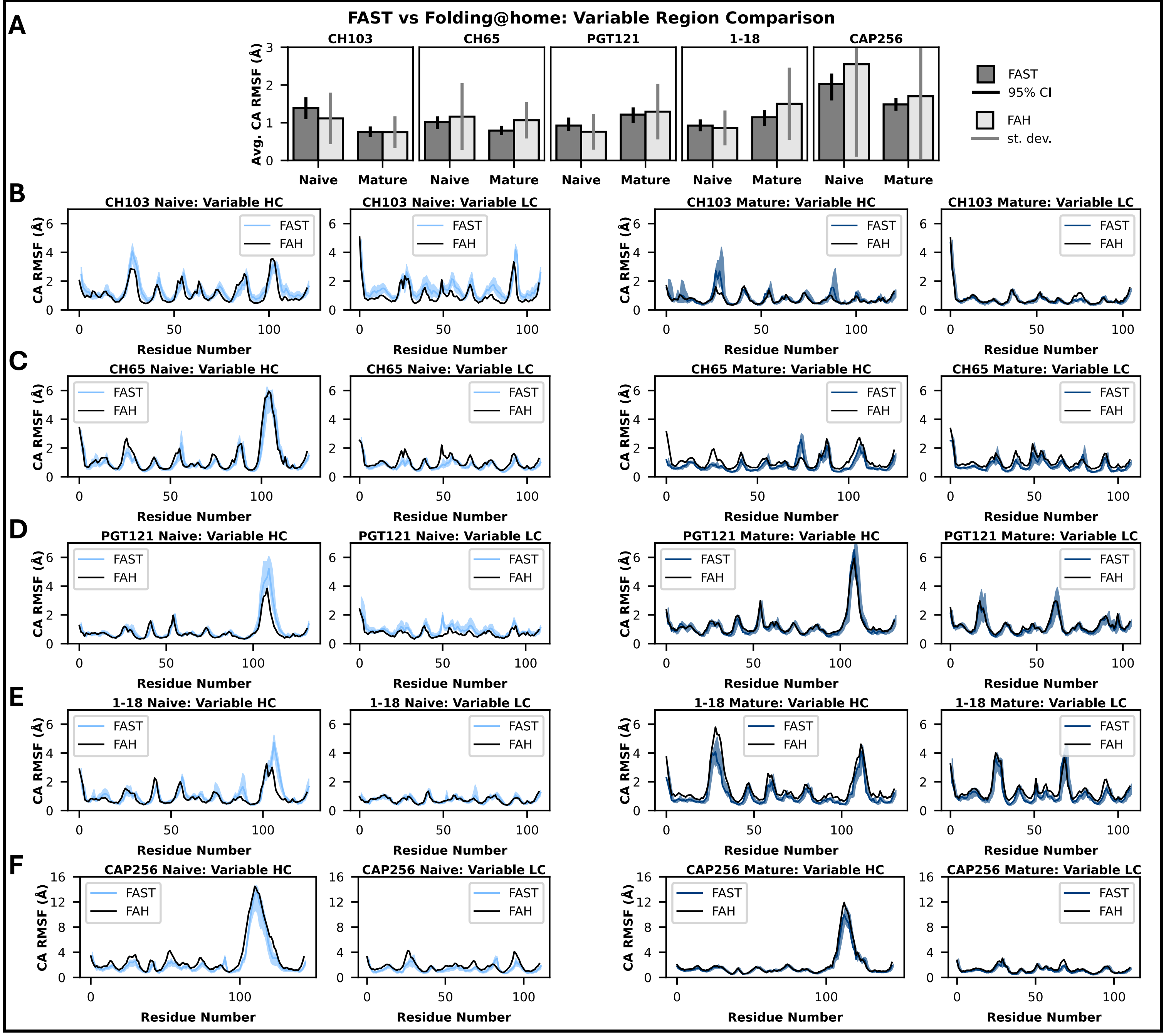


**Figure S1.** Comparison of RMSF values from FAST and Folding@home simulations shows strong agreement.

(A) Averaged RMSF values (of variable heavy and light chains) for FAST (dark gray) and Folding@home (light gray) simulations. The error bars on the FAST data represent a 95% bootstrapped confidence interval. The error bars on the Folding@home (FAH) data represent one standard deviation of the mean; a 95% bootstrapped confidence interval is difficult to obtain due to the large size of the dataset. The average RMSF from Folding@home falls within the 95% confidence interval from the FAST RMSF data in 5 of the 10 antibody simulation sets. The FAST confidence intervals fall within the standard deviation ranges of the Folding@home data in all 10 sets of simulations.

(B) through (F): Per-residue RMSF profile plots comparing FAST (light blue for naïve antibodies, dark blue for mature antibodies) vs Folding@home (black) simulations for lineages CH103, CH65, PGT121, 1-18, and CAP256, respectively. The shaded regions surrounding the FAST RMSF profiles represent a 95% bootstrapped confidence interval. The RMSF profiles show high similarity, with a majority of the per-residue RMSF values from Folding@home falling within the 95% confidence intervals from the FAST data.

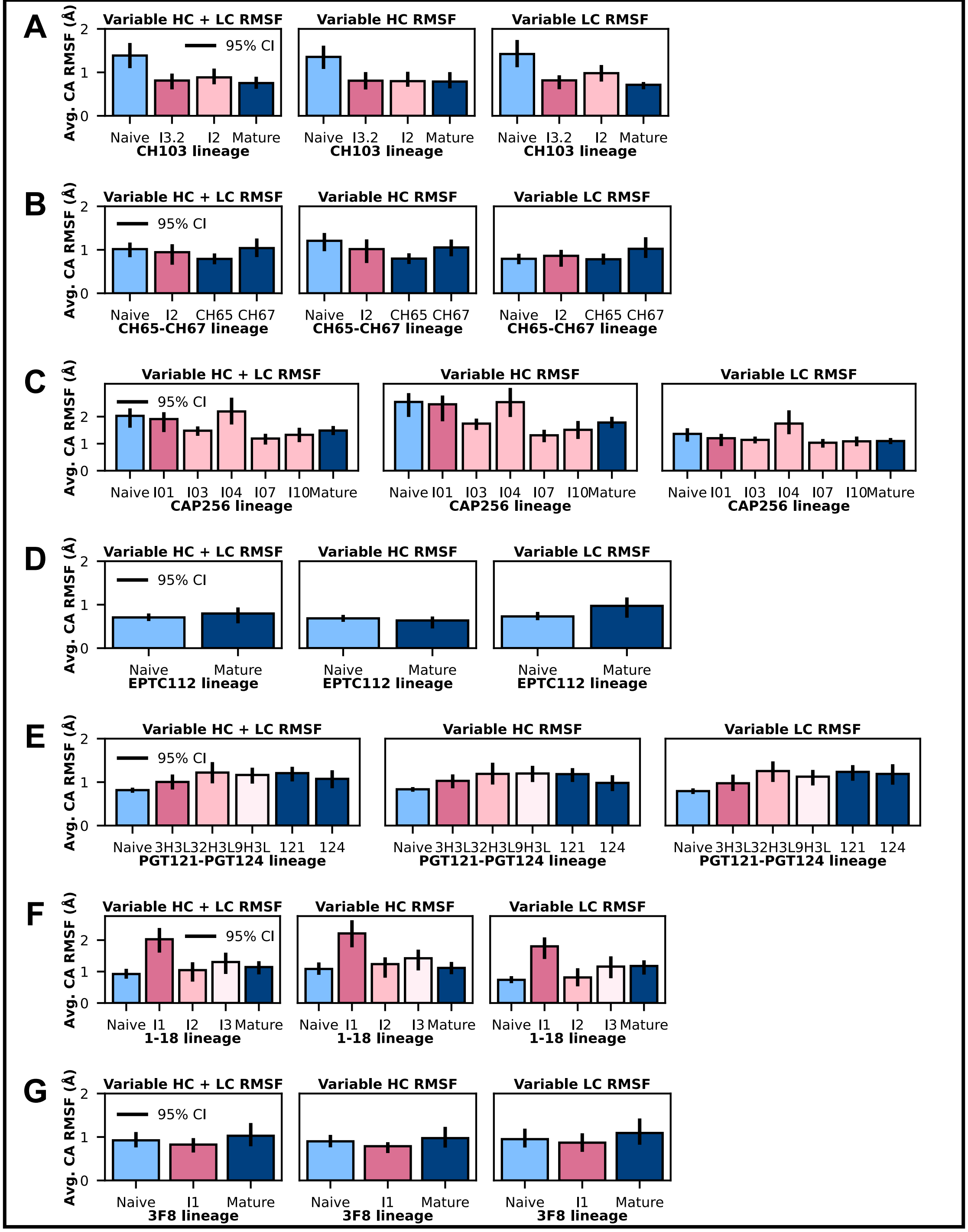


**Figure S2.** There is not a consistent trend in flexibility with maturation across all lineages.
(A) - (G) RMSF plots for averaged variable region (HC + LC), variable heavy chain (HC), and variable light chain (LC) with bootstrapped 95% confidence intervals as error bars for each antibody. The naïve antibody is in light blue, mature antibodies are in dark blue, and any intermediates are in shades of pink.

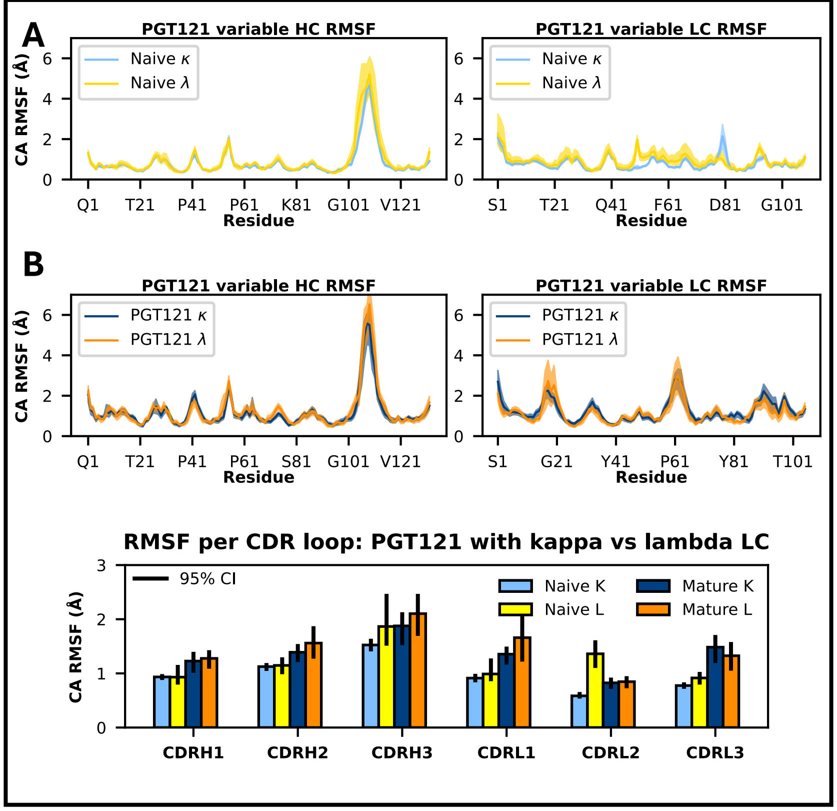


**Figure S3.** Comparison of variable region RMSF from swapping light chain constants in PGT121 shows strong agreement.

(A) Per-residue RMSF values for the variable heavy and light chains for naïve PGT121 with the kappa (light blue) vs with lambda (yellow) light chain constant show general agreement, with disagreements at two regions, around residues 50 and 81. The shaded region represents a bootstrapped 95% confidence interval.

(B) Per-residue RMSF values for the variable heavy and light chains for mature PGT121 with the kappa (dark blue) vs with lambda (orange) light chain constant show strong agreement. The shaded region represents a bootstrapped 95% confidence interval.

(C) Averaged RMSF values for each CDR loop for each naïve/mature and light chain constant combination shows agreement in 11 of the 12 comparisons. The one comparison that does not have overlapping 95% confidence intervals is the CDRL2 of the naïve antibodies. The error bars represent bootstrapped 95% confidence intervals.

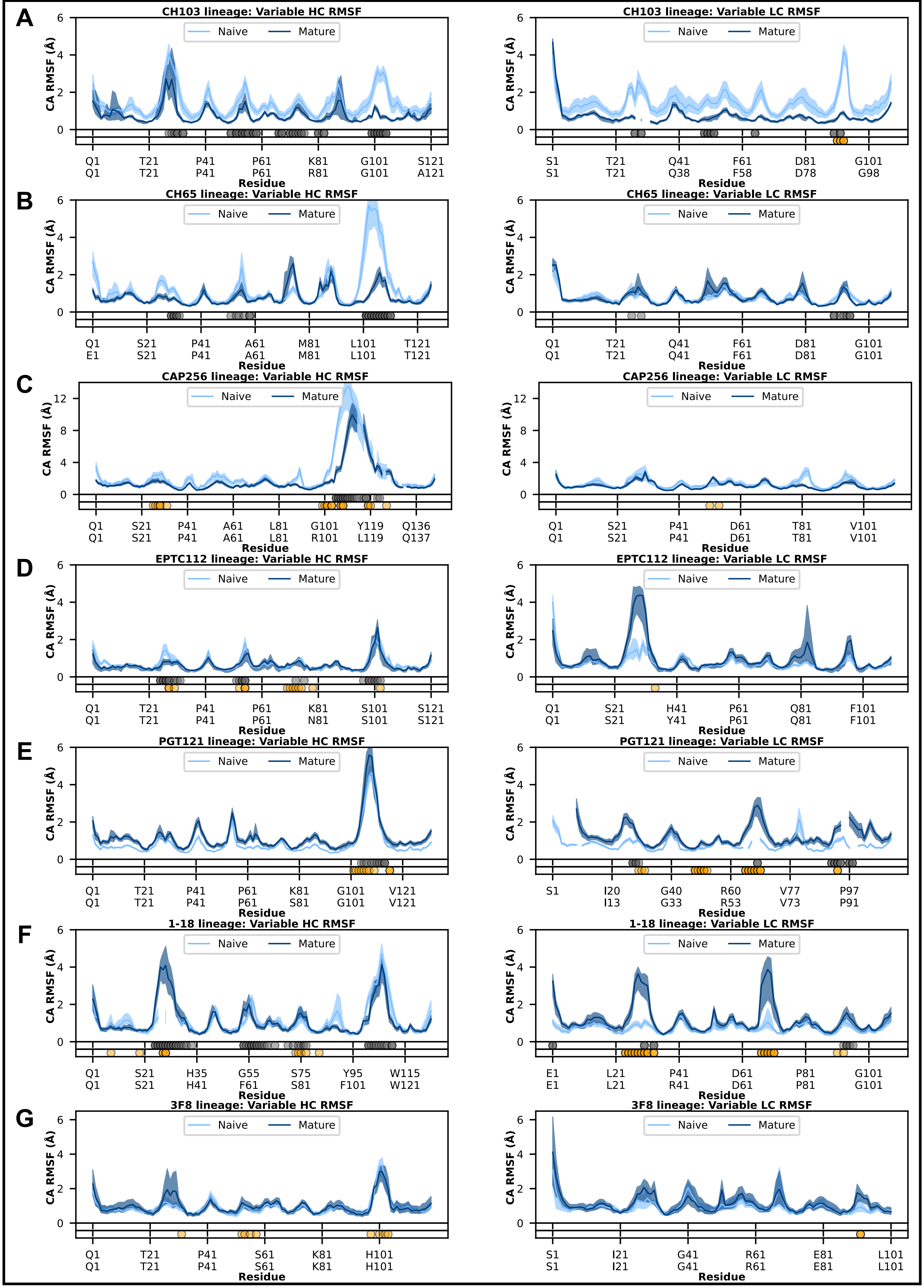


**Figure S4.** Sequence-aligned RMSF plots with shaded bootstrapped 95% confidence intervals for all naïve and mature antibodies. Antibody residues within 6 Å of an antigen protein residue or glycan in the corresponding mature PDB structure are denoted by gray and orange dots, respectively, under the RMSF plot. Opacity of the dot indicates a significant difference in RMSF between the naïve and mature antibody at that residue. (A) – (G): Lineages CH103, CH65, CAP256, EPTC112, PGT121, 1-18, and 3F8, respectively.

A

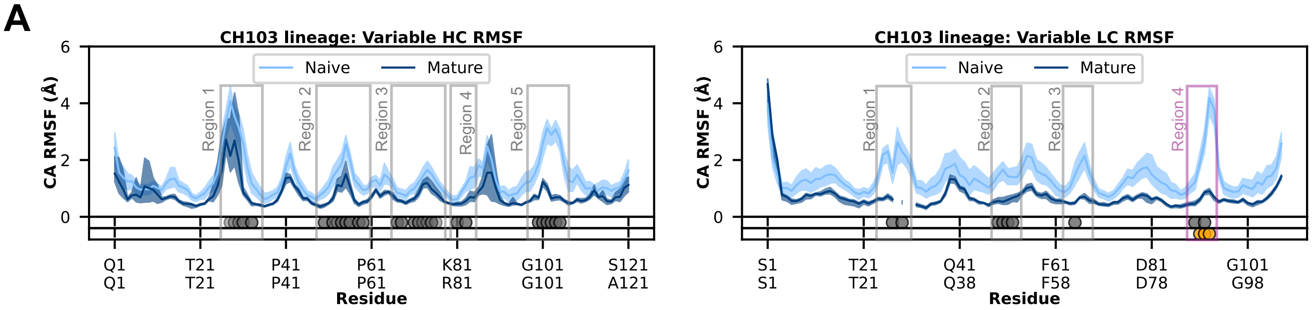


B

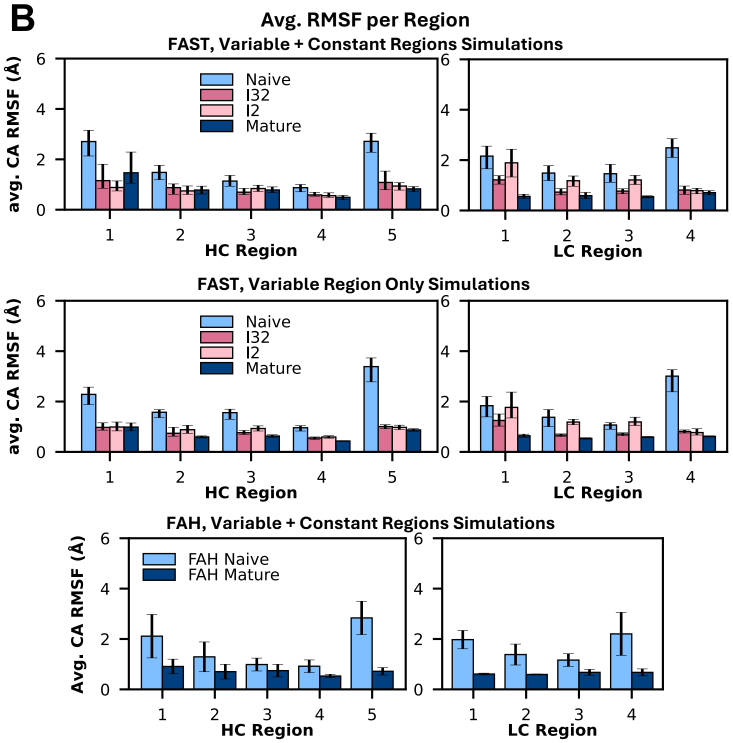


C

**Residues in each region**

| Antibody | HC Region 1 | HC Region 2 | HC Region 3 | HC Region 4 | HC Region 5 |
|---|---|---|---|---|---|
| **Naïve** | SISSYY | YIYYSGSTNY | VTISVDTSKN | KLS | RGQLVN |
| **I3.2** | SMGGYY | YIFHTGHTNY | VTVSVDTSEN | RLR | RGQLVN |
| **I2** | SMGGTY | YIFHTGHTNY | VSISVDTSED | RLR | RGQLVN |
| **Mature** | SMGGTY | YIFHTGETNY | VSISVDTSED | RLR | RGQLVN |

| Antibody | LC Region 1 | LC Region 2 | LC Region 3 | LC Region 4 |
|---|---|---|---|---|
| **Naïve** | LGD | QDSK | SNS | WDSF |
| **I3.2** | LGD | QDSK | SNS | WDSF |
| **I2** | VAS | ENYK | SKS | WDSF |
| **Mature** | TN | ENYK | SKS | WDSF |

**Figure S5.** CH103 lineage RMSF per region.

(A) Sequence-aligned per-residue RMSF plots for heavy and light chains of the naïve and mature antibodies with bootstrapped 95% confidence intervals shaded. Antibody residues within 6 Å of an antigen protein residue or glycan in the corresponding mature PDB structure are denoted by gray and orange dots, respectively, under the RMSF plot. Opacity of the dot indicates a significant difference in RMSF between the naïve and mature antibody at that residue.

(B) Averaged RMSF values per region in (A) across all lineage members, with bootstrapped 95% confidence intervals shown as error bars. The first plot is from the regular simulations (FAST, variable + constant regions). The second plot is from the FAST variable-only simulations. The third plot is from the Folding@home simulations.

(C) Residues included in each region's RMSF averaging in (B). If a region only contained one residue, then one surrounding residue was also included, making the region three residues long.

A

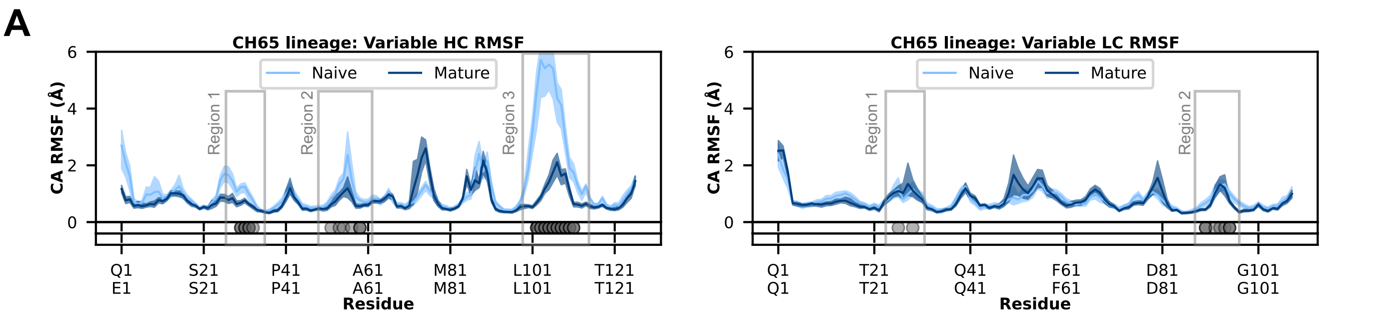


B

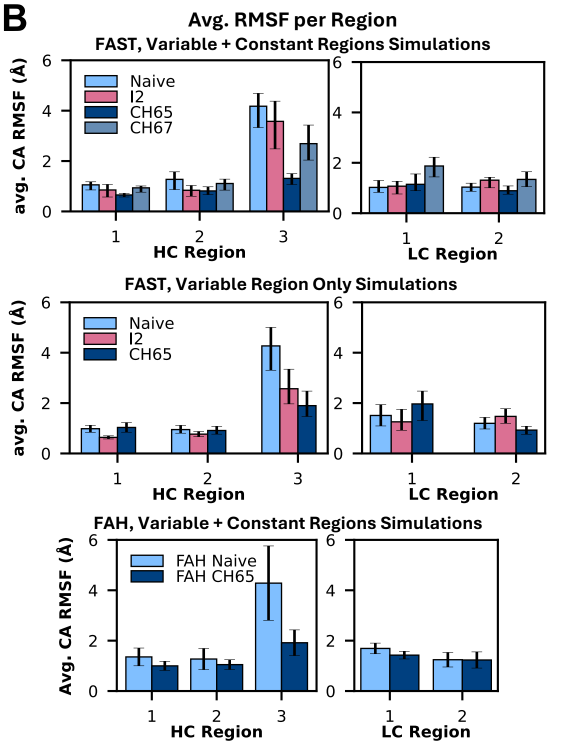


C

Residues in each region

| Antibody | HC Region 1 | HC Region 2 | HC Region 3 | LC Region 1 | LC Region 2 |
|---|---|---|---|---|---|
| Naïve | TGYY | NPNSGGTN | EPRSVDYYYY | NIGS | WDSSSD |
| I2 | TDYY | HPNSGGTN | EPRSVDYYYY | NIGS | WDSSSD |
| CH65 Mature | TDYH | HPNSGDTN | EPRSVDYYYY | DIGR | WDSSSD |
| CH67 Mature | TDNY | HPNSGATK | EPRSVDYYFY | NIGR | WDSDSD |

**Figure S6.** CH65 lineage RMSF per region.

(A) Sequence-aligned per-residue RMSF plots for heavy and light chains of the naïve and mature antibodies with bootstrapped 95% confidence intervals shaded. Antibody residues within 6 Å of an antigen protein residue or glycan in the corresponding mature PDB structure are denoted by gray and orange dots, respectively, under the RMSF plot. Opacity of the dot indicates a significant difference in RMSF between the naïve and mature antibody at that residue.

(B) Averaged RMSF values per region in (A) across all lineage members, with bootstrapped 95% confidence intervals shown as error bars. The first plot is from the regular simulations (FAST, variable + constant regions). The second plot is from the FAST variable-only simulations. The third plot is from the Folding@home simulations.

(C) Residues included in each region's RMSF averaging in (B).

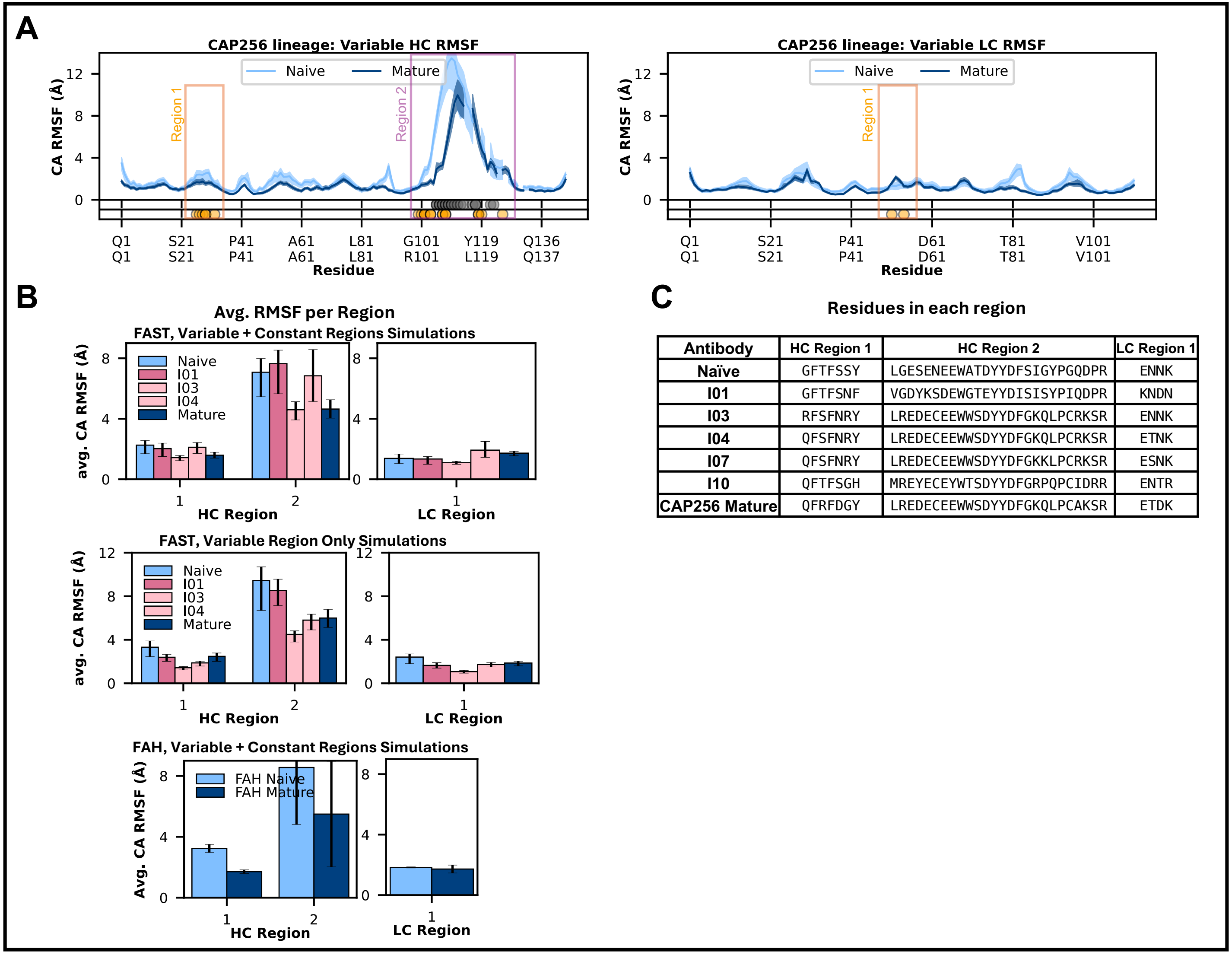


| Antibody | HC Region 1 | HC Region 2 | LC Region 1 |
|---|---|---|---|
| **Naïve** | GFTFSSY | LGESENEEWATDYYDFSIGYPGQDPR | ENNK |
| **I01** | GFTFSNF | VGDYKSDEWGTEYYDISISYPIQDPR | KNDN |
| **I03** | RFSFNRY | LREDECEEWWSDYYDFGKQLPCRKSR | ENNK |
| **I04** | QFSFNRY | LREDECEEWWSDYYDFGKQLPCRKSR | ETNK |
| **I07** | QFSFNRY | LREDECEEWWSDYYDFGKKLPCRKSR | ESNK |
| **I10** | QFTFSGH | MREYECEYWTSDYYDFGRPQPCIDRR | ENTR |
| **CAP256 Mature** | QFRFDGY | LREDECEEWWSDYYDFGKQLPCAKSR | ETDK |

**Figure S7.** CAP256 lineage RMSF per region.

(A) Sequence-aligned per-residue RMSF plots for heavy and light chains of the naïve and mature antibodies with bootstrapped 95% confidence intervals shaded. Antibody residues within 6 Å of an antigen protein residue or glycan in the corresponding mature PDB structure are denoted by gray and orange dots, respectively, under the RMSF plot. Opacity of the dot indicates a significant difference in RMSF between the naïve and mature antibody at that residue.

(B) Averaged RMSF values per region in (A) across all lineage members, with bootstrapped 95% confidence intervals shown as error bars. The first plot is from the regular simulations (FAST, variable + constant regions). The second plot is from the FAST variable-only simulations. The third plot is from the Folding@home simulations.

(C) Residues included in each region's RMSF averaging in (B).

A

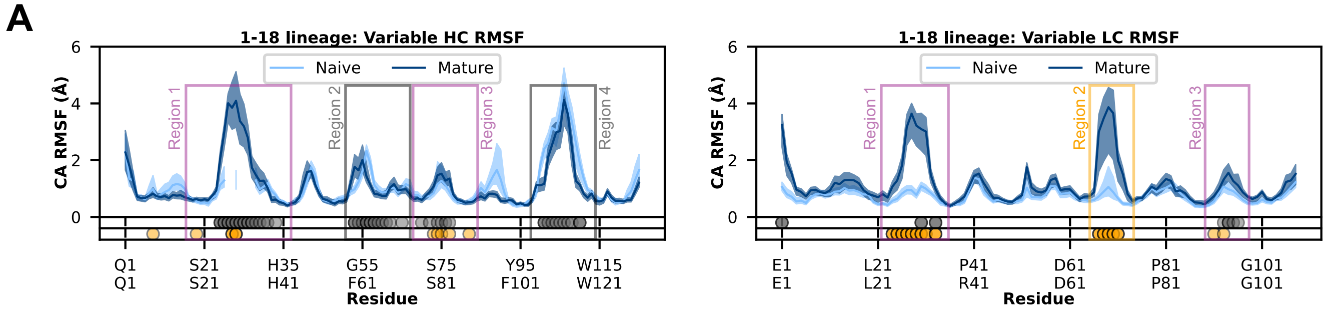


B

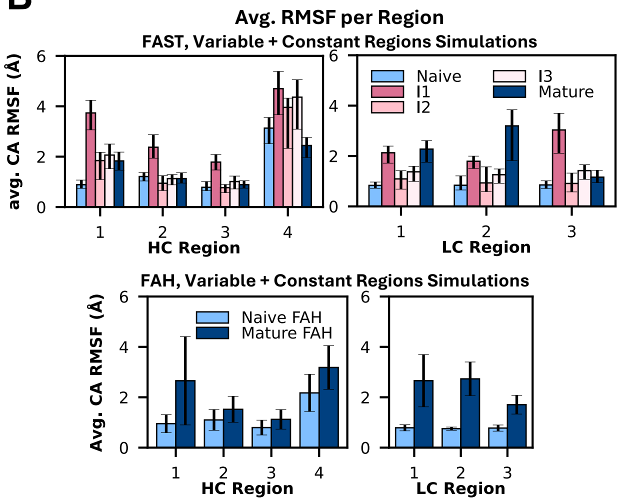


C

**Residues in each region**

| Antibody | HC Region 1 | HC Region 2 | HC Region 3 | HC Region 4 |
|---|---|---|---|---|
| Naïve | KVSCKASGYTFTSYY | PSGGSTSYAQKFQ | MTRDTSTSTVYME | FGERAPYYYY |
| I1 | KVSCRADDDPYTDDDTFTKYW | PHFARPIYSYKFRGR | MTRDTSLTTVYMEL | FGDRAPHYNY |
| I2 | KVSCRADDDPYTDDDTFTKYW | PHFARPIYSYKFRDR | MTRDTSLTAVYMEL | FGDRAPHYNY |
| I3 | RISCRADDDPYTDDDTFTKYY | PHFARPIYSYKFQDR | LTRDSSLTAVYLEL | FGERAPHYNY |
| 1-18 Mature | RISCRADDDPYTDDDTFTKYW | PHFARPIYSYKFRDR | LTRDSSLTAVYLEL | FGDRAPHYNY |

| Antibody | LC Region 1 | LC Region 2 | LC Region 3 |
|---|---|---|---|
| Naïve | RASQSVSSSY | GSGTD | QYGSSP |
| I1 | RASQGLSSSH | GSGTD | RYGSTP |
| I2 | RASQGLSSSH | GSGTA | RYGNTP |
| I3 | RASEGLDSSQ | GSGAD | RFGATP |
| 1-18 Mature | RASQGLDSSH | GSGAD | RYGGTP |

**Figure S8.** 1-18 lineage RMSF per region.

(A) Sequence-aligned per-residue RMSF plots for heavy and light chains of the naïve and mature antibodies with bootstrapped 95% confidence intervals shaded. Antibody residues within 6 Å of an antigen protein residue or glycan in the corresponding mature PDB structure are denoted by gray and orange dots, respectively, under the RMSF plot. Opacity of the dot indicates a significant difference in RMSF between the naïve and mature antibody at that residue.

(B) Averaged RMSF values per region in (A) across all lineage members, with bootstrapped 95% confidence intervals shown as error bars. The first plot is from the regular simulations (FAST, variable + constant regions). The second plot is from the Folding@home simulations.

(C) Residues included in each region's RMSF averaging in (B).

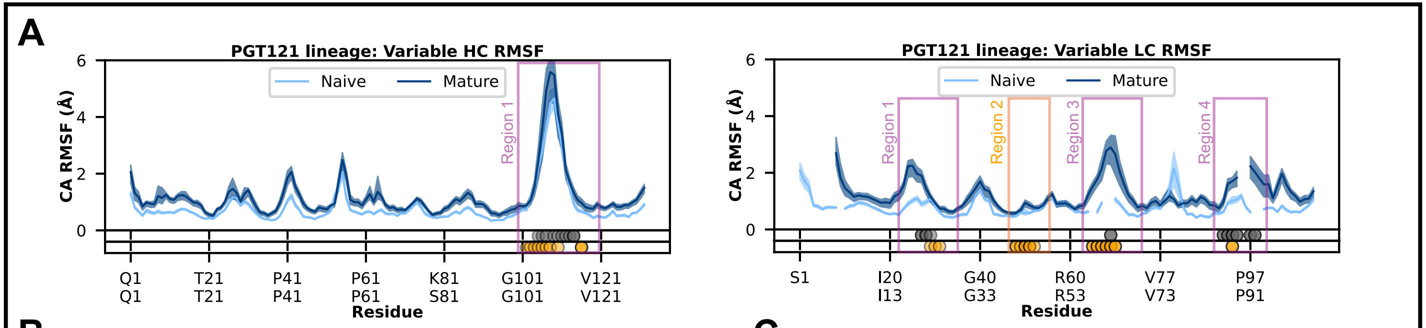


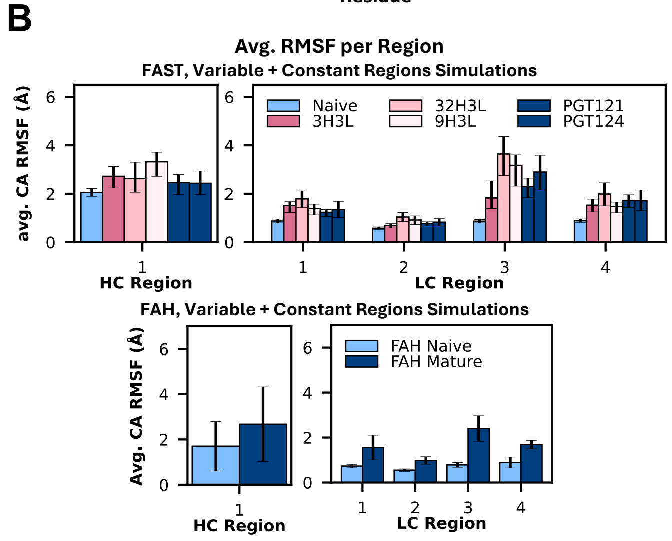


C

**Residues in each region**

| Antibody | HC Region 1 | LC Region 1 | LC Region 2 | LC Region 3 | LC Region 4 |
|---|---|---|---|---|---|
| Naïve | KRIYGVVSFGDYYYYYY | IGSKS | YDDSD | NSG | DSSSDHP |
| 3H3L | KRIYGMVSFGEFFYYYY | LGSRA | YNNQD | PDSNFG | DSRSAIN |
| 32H3L | QRIYGVVSFGEFFYYYY | LGSRA | YNNQD | PDSNFG | DSRSAIN |
| 9H3L | RRIYGIVAFREWFTYFY | LGSRA | YNNQD | PDSNFG | DSRSAIN |
| PGT121 | RRIYGIVAFNEWFTYFY | LGSRA | YNNQD | PDSPFG | DSRVPT |
| PGT124 | QRIYGVVSFGEFFYYYY | LGSRA | YNNQD | PDINFG | DSRSGFS |

**Figure S9.** PGT121 lineage RMSF per region.

(A) Sequence-aligned per-residue RMSF plots for heavy and light chains of the naïve and mature antibodies with bootstrapped 95% confidence intervals shaded. Antibody residues within 6 Å of an antigen protein residue or glycan in the corresponding mature PDB structure are denoted by gray and orange dots, respectively, under the RMSF plot. Opacity of the dot indicates a significant difference in RMSF between the naïve and mature antibody at that residue.

(B) Averaged RMSF values per region in (A) across all lineage members, with bootstrapped 95% confidence intervals shown as error bars. The first plot is from the regular simulations (FAST, variable + constant regions). The second plot is from the Folding@home simulations.

(C) Residues included in each region's RMSF averaging in (B).

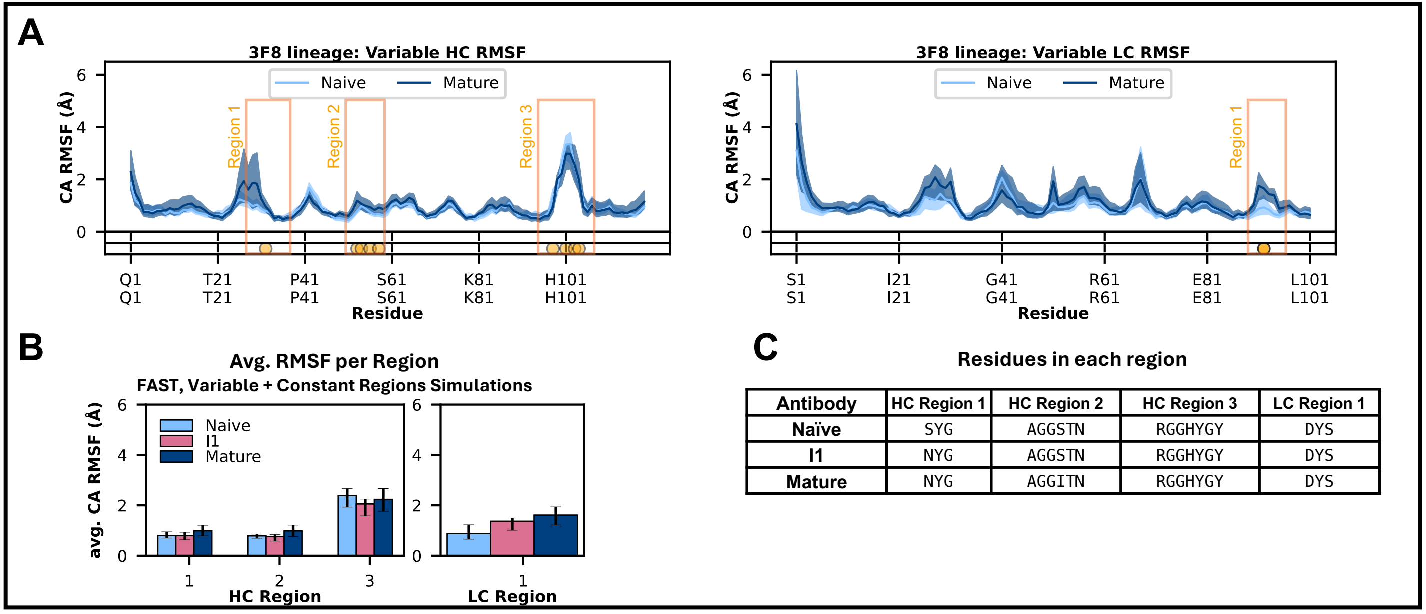


| Antibody | HC Region 1 | HC Region 2 | HC Region 3 | LC Region 1 |
|---|---|---|---|---|
| Naïve | SYG | AGGSTN | RGGHYGY | DYS |
| I1 | NYG | AGGSTN | RGGHYGY | DYS |
| Mature | NYG | AGGITN | RGGHYGY | DYS |

**Figure S10.** 3F8 lineage RMSF per region.

(A) Sequence-aligned per-residue RMSF plots for heavy and light chains of the naïve and mature antibodies with bootstrapped 95% confidence intervals shaded. Antibody residues within 6 Å of an antigen protein residue or glycan in the corresponding mature PDB structure are denoted by gray and orange dots, respectively, under the RMSF plot. Opacity of the dot indicates a significant difference in RMSF between the naïve and mature antibody at that residue.

(B) Averaged RMSF values per region in (A) across all lineage members, with bootstrapped 95% confidence intervals shown as error bars; the data is from the regular simulations (FAST, variable + constant regions).

(C) Residues included in each region's RMSF averaging in (B). If a region only contained one residue, then one surrounding residue was also included, making the region three residues long.

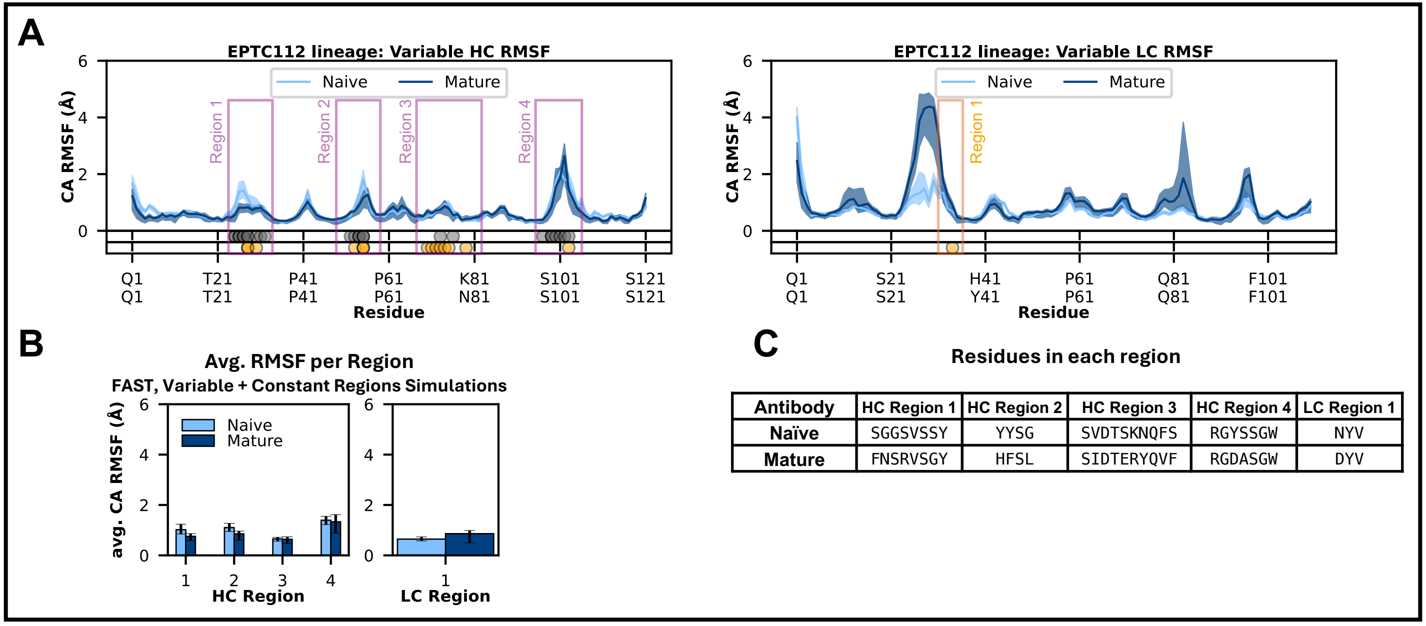


| Antibody | HC Region 1 | HC Region 2 | HC Region 3 | HC Region 4 | LC Region 1 |
|---|---|---|---|---|---|
| Naïve | SGGSVSSY | YYSG | SVDTSKNQFS | RGYSSGW | NYV |
| Mature | FNSRVSGY | HFSL | SIDTERYQVF | RGDASGW | DYV |

**Figure S11.** EPTC112 lineage RMSF per region.

(A) Sequence-aligned per-residue RMSF plots for heavy and light chains of the naïve and mature antibodies with bootstrapped 95% confidence intervals shaded. Antibody residues within 6 Å of an antigen protein residue or glycan in the corresponding mature PDB structure are denoted by gray and orange dots, respectively, under the RMSF plot. Opacity of the dot indicates a significant difference in RMSF between the naïve and mature antibody at that residue.

(B) Averaged RMSF values per region in (A) across all lineage members, with bootstrapped 95% confidence intervals shown as error bars; the data is from the regular simulations (FAST, variable + constant regions).

(C) Residues included in each region's RMSF averaging in (B). If a region only contained one residue, then one surrounding residue was also included, making the region three residues long.

**Table S1.** Sequence information for each antibody simulated. The numbering for the intermediates is not necessarily in order of when they were isolated or their neutralization strength.

| Lineage Name | Antibody Name | Source and Description |
|---|---|---|
| 1-18 | Naïve | Inferred germline sequence. Designed using the predicted VDJ and VJ genes of the mature 1-18 sequence. |
| 1-18 | I1 | Combinatorial analysis of VH1-46 class antibodies isolated from individual IDC561 (human). |
| 1-18 | I2 | Isolated after immunization with the native HIV trimer in an immunoglobulin knock-in mouse expressing the I1 antibody. |
| 1-18 | I3 | Schommers et al. 2020, *Cell* \| Antibody 561_09_23. Isolated from individual IDC561 (human). This was chosen as an intermediate because it comes from the same clone as the mature 1-18 antibody (Clone 4.1), has weaker potency and neutralization compared to 1-18, and is closer in sequence identity to the naïve antibody. |
| 1-18 | Mature | Schommers et al. 2020, *Cell* \| Antibody 561_01_18. Mature antibody from 1-18 lineage. Isolated from individual IDC561 (human), Clone 4.1. |
| CH65-CH67 | Naïve | Whittle et al. 2011, *PNAS* \| Unmutated common ancestor. \| PBD ID: 4HK0 |
| CH65-CH67 | I2 | Whittle et al. 2011, *PNAS* \| Isolated from human subject after influenza vaccination. CH65 and CH67 diverge into two branches at this intermediate sequence. \| PDB ID: 4HK3 |
| CH65-CH67 | CH65 Mature | Whittle et al. 2011, *PNAS* \| Isolated from human subject after influenza vaccination. \| PDB ID: 5UGY |
| CH65-CH67 | CH67 Mature | Whittle et al. 2011, *PNAS* \| Isolated from human subject after influenza vaccination. |
| 3F8 | Naïve | Sterner et al. 2017, *Cell* \| Inferred germline sequence. |
| 3F8 | I1 | Sterner et al. 2017, *Cell* \| Germline reversion (I56S) of Mature 3F8. |
| 3F8 | Mature | Sterner et al. 2017, *Cell* \| Murine monoclonal antibody. |
| CH103 | Naïve | Liao et al. 2013, *Nature* \| Unmutated common ancestor. \| PDB ID: 4QHK |
| CH103 | I3.2 | Liao et al. 2013, *Nature.* Fera et al. 2014, *PNAS.* \| Isolated from donor CH505 (human). I3.2 is the I3 heavy chain with the I2 light chain. \| PDB ID: 4QHL |
| CH103 | I2 | Liao et al. 2013, *Nature* \| Isolated from donor CH505 (human). \| PDB ID: 4QHN |
| CH103 | Mature | Liao et al. 2013, *Nature* \| Isolated from donor CH505 (human). \| PDB ID: 4JAM |
| EPTC112 | Naïve | Inferred germline sequence (inferred using sequences reported in Molinos-Albert et al. 2023, *Cell Host and Microbe).* |
| EPTC112 | Mature | Molinos-Albert et al. 2023, *Cell Host and Microbe* \| Isolated from a post-treatment controller individual (human). |
| PGT121-PGT124 | Naïve | Garces et al. 2014, *Cell* \| Inferred germline sequence. |
| PGT121-PGT124 | I1 | Garces et al. 2014, *Cell* \| Isolated from African donor 17 of the IAVI Protocol G cohort (human). 3H + 3L. Precursor for PGT121, 32H + 3L, and 9H + 3L. |
| PGT121-PGT124 | I2 | Garces et al. 2014, *Cell* \| Isolated from African donor 17 of the IAVI Protocol G cohort (human). 32H + 3L. Precursor for PGT124. |
| PGT121-PGT124 | I3 | Garces et al. 2014, *Cell* \| Isolated from African donor 17 of the IAVI Protocol G cohort (human). 9H + 3L. Precursor for PGT122 (PGT122 was not simulated in this study). |
| PGT121-PGT124 | PGT121 Mature | Garces et al. 2014, *Cell* \| Isolated from African donor 17 of the IAVI Protocol G cohort (human). |
| PGT121-PGT124 | PGT124 Mature | Garces et al. 2014, *Cell* \| Isolated from African donor 17 of the IAVI Protocol G cohort (human). |
| CAP256 | Naïve | Doria-Rose et al. 2014, *Nature* \| Unmutated common ancestor. \| PDB ID: 4ODH |
| CAP256 | I01 | Doria-Rose et al. 2014, *Nature* \| CAP256-VRC26.01, isolated at week 59 post-infection from donor CAP256 (human). \| PDB ID: 4ORC |
| CAP256 | I03 | Doria-Rose et al. 2014, *Nature* \| CAP256-VRC26.03, isolated at week 119 post-infection from donor CAP256 (human). \| PDB ID: 4OD1 |
| CAP256 | I04 | Doria-Rose et al. 2014, *Nature* \| CAP256-VRC26.04, isolated at week 119 post-infection from donor CAP256 (human). \| PDB ID: 4ORG |
| CAP256 | I07 | Doria-Rose et al. 2014, *Nature* \| CAP256-VRC26.07, isolated at week 119 post-infection from donor CAP256 (human). \| PDB ID: 4OD3 |
| CAP256 | I10 | Doria-Rose et al. 2014, *Nature* \| CAP256-VRC26.10, isolated at week 206 post-infection from donor CAP256 (human). \| PDB ID: 4OCS |
| CAP256 | Mature | Doria-Rose et al. 2015, *Journal of virology* \| CAP256-VRC26.25, isolated at week 193 post-infection from donor CAP256 (human). \| PDB ID: 5DT1 |

**Table S2.** Variable region sequences for all antibodies simulated. The CDR1-3 loops are highlighted in each naïve and mature antibody according to the IMGT definition scheme (yellow for CDRH loops, orange for CDRL loops).

| HC/ LC | Antibody | Variable Region Sequence |
|---|---|---|
| HC | 1-18 Naive | QVQLVQSGAEVKKPGASVKVSCKASGYTFTSYYMHWVRQAPGQGLEWMGIINPSGGSTSYAQKFQGRVTMTRDTSTSTVYMELSSLRSEDTAVYYCARDPFGERAPYYYYYMDVWGGGTAVIVSS |
| | 1-18 I1 | QVQLVQSGAEVKKPGASVKVSCRADDDPYTDDDTFTKYWMHWIRQAPGQRPEWLGVISPHFARPIYSYKFRGRVTMTRDTSLTTVYMELSSLRSEDSGVYYCARDPFGDRAPHYNYHMDVWGGGTAVIVSS |
| | 1-18 I2 | QEQLVQSGAEVKKPGASVKVSCRADDDPYTDDDTFTKYWTHWIRQAPGQRPEWLGVISPHFARPIYSYKFRDRLTMTRDTSLTAVYMELSGLSSEDSGVYYCARDPFGDRAPHYNYHMDVWGGGTAVIVSS |
| | 1-18 I3 | QAHLFQSGAELKRPGASVRISCRADDDPYTDDDTFTKYYTHWIRQAPGQRPEWLGVISPHFARPIYSYKFQDRLTLTRDSSLTAVYLELRSLRLDDTGIYYCARDPFGERAPHYNYHMDVWGAGTTVIVSS |
| | 1-18 Mature | QGRLFQSGAEVKRPGASVRISCRADDDPYTDDDTFTKYWTHWIRQAPGQRPEWLGVISPHFARPIYSYKFRDRLTLTRDSSLTAVYLELKGLQPDDSGIYFCARDPFGDRAPHYNYHMDVWGGGTAVIVSS |
| LC | 1-18 Naive | EIVLTQSPGTLSLSPGERATLSCRASQSVSSSYLAWYQQKPGQAPRLLIYGASSRATGIPDRFSGSGSGTDFTLTISRLEPEDFAVYYCQQYGSSPLTFGGGTKVEIK |
| | 1-18 I1 | EVVLTQSPGTLSLSPGERATLSCRASQGLSSSHLAWYQQKPGQAPRLLIFGTSNRARGIPDRFSGSGSGTDFTLTISRVEPEDFAVYYCQRYGSTPITFGGGTTLDKK |
| | 1-18 I2 | EVVLTQSPGTLSLSPGERATLSCRASQGLSSSHLAWYQQKPGQAPRLLIFGTSNRARGIPDRFSGSGSGTAFTLTISRVEPEDFAVYYCQRYGNTPITFGGGTTLDKK |
| | 1-18 I3 | EVVLTQSPAILSASPGDRVVLSCRASEGLDSSQLAWYRFKDGQIPRLVLFGVSNRARGTPDRFSGGGSGADFTLTISRVEREDFATYYCQRFGATPITFGGGTRLDMN |
| | 1-18 Mature | EVVLTQSPAILSVSPGDRVILSCRASQGLDSSHLAWYRFKRGQIPTLVIFGTSNRARGTPDRFSGSGSGADFTLTISRVEPEDFATYYCQRYGGTPITFGGGTTLDKK |
| HC | CH65 Naive | QVQLVQSGAEVKKPGASVKVSCKASGYTFTGYYMHWVRQAPGQGLEWMGWINPNSGGTNYAQKFQGWVTMTRDTSISTAYMELSRLRSDDTAVYYCARGGLEPRSVDYYYYGMDVWGQGTTVTVSS |
| | CH65 I2 | QVQLVQSGAEVKKPGASVKVSCKASGYTFTDYYIHWVRQAPGQGLEWMGWIHPNSGGTNYAQKFQGWVTMTRDTSISTAYMELSRLRSDDTAVYYCARGGLEPRSVDYYYYGMDVWGQGTTVTVSS |
| | CH65 Mat. | EVQLVQSGAEVKKPGASVKVSCKASGYTFTDYHINWVRQAPGQGLEWMGWIHPNSGDTNYAQKFQGWVTMTRDTAISTAYMEVNGLKSDDTAVYYCARGGLEPRSVDYYYYGMDVWGQGTTVTVSS |
| | CH67 Mat. | QVQLVQSGAEVRKPGASVKVSCKASGYTFTDNYIHWVRQAPGQGLEWMGWIHPNSGATKYAQKFEGWVTMTRDTSISTVYMELSRSRSDDTAVYYCARAGLEPRSVDYYFYGLDVWGQGTAVTVSS |
| LC | CH65 Naive | QSVLTQPPSVSVAPGQTARITCGGNNIGSKSVHWYQQKPGQAPVLVVYDDSDRPSGIPERFSGSNSGNTATLTISRVEAGDEADYYCQVWDSSSDHVVFGGGTKLTVL |
| | CH65 I2 | QSVLTQPPSVSVAPGQTARITCGGNNIGSKSVHWYQQKPGQAPVLVVYDDSDRPSGIPERFSGSNSGNTATLTISRVEAGDEADYYCQVWDSSSDHVVFGGGTKLTVL |
| | CH65 Mat. | QSVLTQPPSVSVAPGQTARITCGGNDIGRKSVHWNQQKPGQAPVLVVCYDSDRPSGIPERFSGSNSGNTATLTISRVEAGDEADYYCQVWDSSSDHVIFGGGTKLTVL |
| | CH67 Mat. | QSALTQPPSVSVAPGQTATITCGGNNIGRKRVDWFQQKPGQAPVLVVYEDSDRPSGIPERFSDSNSGTTATLTISRVEAGDEADYYCQVWDSDSDHVVFGGGTKLTVL |
| HC | 3F8 Naive | QVQLKESGPGLVAPSQSLSITCTVSGFSLTSYGVHWVRQPPGKGLEWLGVIWAGGSTNYNSALMSRLSISKDNSKSQVFLKMNSLQTDDTAMYYCASRGGHYGYAMDYWGQGTSVTVSS |
| | 3F8 I1 | QVQLKESGPGLVAPSQSLSITCTVSGFSVTNYGVHWVRQPPGKGLEWLGVIWAGGSTNYNSAFMSRLSISKDNSKSQVFLKMNSLQIDDTAMYYCASRGGHYGYALDYWGQGTSVTVSS |
| | 3F8 Mature | QVQLKESGPGLVAPSQSLSITCTVSGFSVTNYGVHWVRQPPGKGLEWLGVIWAGGITNYNSAFMSRLSISKDNSKSQVFLKMNSLQIDDTAMYYCASRGGHYGYALDYWGQGTSVTVSS |
| LC | 3F8 Naive | SIVMTQTPKFLLVSAGDRVTITCKASQSVSNDVAWYQQKAGQSAKLLIYYASNRYTGVPDRFTGSGYGTDFTFTISTVQAEDLAVYFCQQDYSSFGGGTKL |
| | 3F8 I1 | SIVMTQTPKFLLVSAGDRVTITCKASQSVSNDVTWYQQKAGQSPKLLIYSASNRYSGVPDRFTGSGYGTAFTFTISTVQAEDLAVYFCQQDYSSFGGGTKL |
| | 3F8 Mature | SIVMTQTPKFLLVSAGDRVTITCKASQSVSNDVTWYQQKAGQSPKLLIYSASNRYSGVPDRFTGSGYGTAFTFTISTVQAEDLAVYFCQQDYSSFGGGTKL |
| HC | CH103 Naïve | QVQLQESGPGLVKPSETLSLTCTVSGGSISSYYWSWIRQPPGKGLEWIGYIYYSGSTNYNPSLKSRVTISVDTSKNQFSLKLSSVTAADTAVYYCASLPRGQLVNAYFDYWGQGTLVTVSS |
| | CH103 I32 | QVQLQESGPGLVKSSETLSLTCTVSGGSMGGYYWSWLRQSPVKGLEWIGYIFHTGHTNYNPSLESRVTVSVDTSENQFSLRLRSVTAADTAVYYCASLPRGQLVNAFFDNWGQGTLVTVAS |
| | CH103 I2 | QVQLQESGPGVVKSSETLSLTCTVSGGSMGGTYWSWLRLSPGKGLEWIGYIFHTGHTNYNPSLEGRVSISVDTSEDQFSLRLRSVTAADTAVYFCASLPRGQLVNAYFRNWGRGTLVSVTA |
| | CH103 Mature | QVQLQESGPGVVKSSETLSLTCTVSGGSMGGTYWSWLRLSPGKGLEWIGYIFHTGETNYSPSLKGRVSISVDTSEDQFSLRLRSVTAADTAVYFCASLPRGQLVNAYFRNWGRGSLVSVTA |
| LC | CH103 Naïve | SYELTQPPSVSVSPGQTASITCSGDKLGDKYACWYQQKPGQSPVLVIYQDSKRPSGIPERFSGSNSGNTATLTISGTQAMDEADYYCQAWDSFSTFVFGTGTKVTVL |
| | CH103 I32 | YELTQPPSVSVSPGQTASITCSGDKLGDKYACWYQQKPGQSPVLVIYQDSKRPSGIPERFSGSNSGNTATLTISGTQAMDEADYYCQAWDSFSTFVFGTGTKVTVL |
| | CH103 I2 | SYELTQPPSVSVSPGQTATITCSGDKVASKNVCWYQVKPGQSPEVVMYENYKRPSGIPDRFSGSKSGSTATLTIRGTQATDEADYYCQVWDSFSTFVFGSGTQVTVL |
| | CH103 Mature | SYELTQPPSVSVSPGQTATITCSGASTNVCWYQVKPGQSPEVVIFENYKRPSGIPDRFSGSKSGSTATLTIRGTQAIDEADYYCQVWDSFSTFVFGSGTQVTVL |
| HC | EPTC112 Naïve | QVQLQESGPGLVKPSETLSLTCTVSGGSVSSYYWSWIRQPPGKGLEWIGYIYYSGSTNYNPSLKSRVTISVDTSKNQFSLKLSSVTAADTAVYYCARGYSSGWYAEYFQHWGQGTLVTVSS |
| | EPTC112 Mature | QVQLLESGPGLVRPSETLTLTCSVFNSRVSGYYYSWIRQPPGRGLEWIASTHFSLRPSRNPSLLSRVTTSIDTERYQVFLNMRSVTAADTAVYFCARGDASGWRADYFPHWGQGTLVVVSS |
| LC | EPTC112 Naïve | QSALTQPPSASGSPGQSVTISCTGTSSDVGGYNYVSWYQQHPGKAPKLMIYEVSKRPSGVPDRFSGSKSGNTASLTVSGLQAEDEADYYCSSYAGSNNFVFGGGTKLTVL |
| | EPTC112 Mature | QSVLTQPPSASGSPGQSVTISCTGTSSDIGASDYVSWYQQYPGEAPKVIIYDVTKRPSGVPDRFSGSKSGTTASLTVSGLQAEDEADYYCSSDAGRHTLLFGGGTKVTVL |
| HC | PGT121 Naïve | QVQLQESGPGLVKPSETLSLTCTVSGGSISSYYWSWIRQPPGKGLEWIGYIYYSGSTNYNPSLKSRVTISVDTSKNQFSLKLSSVTAADTAVYYCARTQQGKRIYGVVSFGDYYYYYYMDVWGKGTTVTVSS |
| | PGT121 3H3L | QVQLQESGPGLVKPSETLSLTCTVSGGSISNYYWSWIRQSPGKGLEWIGYISDSESTNYNPSLKSRVIISVDTSKNQLSLKLNSVTAADSAIYYCARAQQGKRIYGMVSFGEFFYYYYMDVWGKGTTVTVSS |
| | PGT121 32H3L | QVQLQESGPGLVKPSETLSVTCIVSGGSISNYYWTWIRQSPGKGLEWIGYISDRETTTYNPSLKSRVVISRDTSKNQLSLKLNSVTAADTAIYYCATARRGQRIYGVVSFGEFFYYYYMDVWGKGTAVTVSS |
| | PGT121 9H3L | QVQLQESGPGLVKPSETLSLTCSVSGASISDHYWSWIRQSPGKGLEWIGYVYDSGDTNYNPSLKSRVNLSLDTSKNQVSLSLTAVTAADSAIYYCARTQHGRRIYGIVAFREWFTYFYMDVWGQGTPVTVSS |
| | PGT121 Mature | QMQLQESGPGLVKPSETLSLTCSVSGASISDSYWSWIRRSPGKGLEWIGYVHKSGDTNYSPSLKSRVNLSLDTSKNQVSLSLVAATAADSGKYYCARTLHGRRIYGIVAFNEWFTYFYMDVWGNGTQVTVSS |
| | PGT124 Mature | QVQLQESGPGLVRPSETLSVTCIVSGGSISNYYWTWIRQSPGKGLEWIGYISDRETTTYNPSLNSRAVISRDTSKNQLSLQLRSVTTADTAIYFCATARRGQRIYGVVSFGEFFYYYYMDVWGKGTAVTVSS |
| LC | PGT121 Naïve | SYVLTQPPSVSVAPGQTARITCGGNNIGSKSVHWYQQKPGQAPVLVVYDDSDRPSGIPERFSGSNSGNTATLTISRVEAGDEADYYCQVWDSSSDHPWVFGGGTKLTVL |
| | PGT121 3H3L | SYVLTQPPSLSVAPGETARISCGGRSLGSRAVQWYQQKPGQAPVLVIYNNQDRPSGIPERFSGSPDSNFGTTATLTISRVEAGDEADYYCHMWDSRSAINWVFGGGTKLTVL |
| | PGT121 32H3L | SYVLTQPPSLSVAPGETARISCGGRSLGSRAVQWYQQKPGQAPVLVIYNNQDRPSGIPERFSGSPDSNFGTTATLTISRVEAGDEADYYCHMWDSRSAINWVFGGGTKLTVL |
| | PGT121 9H3L | SYVLTQPPSLSVAPGETARISCGGRSLGSRAVQWYQQKPGQAPVLVIYNNQDRPSGIPERFSGSPDSNFGTTATLTISRVEAGDEADYYCHMWDSRSAINWVFGGGTKLTVL |
| | PGT121 Mature | SDISVAPGETARISCGEKSLGSRAVQWYQHRAGQAPSLIIYNNQDRPSGIPERFSGSPDSPFGTTATLTITSVEAGDEADYYCHIWDSRVPTKWVFGGGTTLTVL |
| | PGT124 Mature | SYVSPLSVALGETARISCGRQALGSRAVQWYQHKPGQAPILLIYNNQDRPSGIPERFSGTPDINFGTTATLTISGVEVGDEADYYCHMWDSRSGFSWSFGGATRLTVL |
| HC | CAP256 Naive | QVQLVESGGGVVQPGRSLRLSCAASGFTFSSYGMHWVRQAPGKGLEWVAVISYDGSNKYYADSVKGRFTISRDNSKNTLYLQMNSLRAEDTAVYYCAKDLGESENEEWATDYYDFSIGYPGQDPRGVVGAFDIWGQGTMVTVSS |
| | CAP256 I01 | EVQVVESGGGVVQPGRSLRLSCTASGFTFSNFAMGWVRQAPGKGLEWVAFISSDGSNKNYGDSVKGRFTISRDNSKNTVFLQMNSLRVEDTALYYCAKDVGDYKSDEWGTEYYDISISYPIQDPRAMVGAFDLWGQGTMVTVSP |
| | CAP256 I03 | EVQLVESGGGVVQPGKSLRLSCAASRFSFNRYGMHWVRQAPGKGLEWVAAISYDGTDKYHADKVWGRFTISRDNSKNTLYLQMNSLRAEDTALYYCAKDLREDECEEWWSDYYDFGKQLPCRKSRGVAGIFDGWGQGTMVTVSS |
| | CAP256 I04 | EVQLVESGGGVVQPGKSLRLSCAASQFSFNRYGMHWVRQAPGKGLEWVAAISYDGTDKYHADKVWGRFTISRDNSKNTLYLQMNSLRAEDTALYYCAKDLREDECEEWWSDYYDFGKQLPCRKSRGVAGIFDKWGQGTMVIVSS |
| | CAP256 I07 | EVQLVESGGGVVQPGRSLRLSCVGSQFSFNRYGMHWVRQAPGKGLEWVAGISFDGTDRYHADNVWGRFTISRDNSKNTLYLQMSSLRAEDTALYYCAKDLREDECEEWWSDYYDFGKKLPCRKSRGVAGVFDKWGQGTMVTVSS |
| | CAP256 I10 | QAILVESGGGVVQPGRSLRLSCAASQFTFSGHGLHWVRQAPGKGLEWVASISFAGTKMDYADSVKGRFAISRDNSKNTLYLQMNSLRVEDTALYYCAKDMREYECEYWTSDYYDFGRPQPCIDRRGVVGIFDMWGQGTMVTVST |
| | CAP256 Mature | QVQLVESGGGVVQPGTSLRLSCAASQFRFDGYGMHWVRQAPGKGLEWVASISHDGIKKYHAEKVWGRFTISRDNSKNTLYLQMNSLRPEDTALYYCAKDLREDECEEWWSDYYDFGKQLPCAKSRGGLVGIADNWGQGTMVTVSS |
| LC | CAP256 Naive | QSVLTQPPSVSAAPGQKVTISCSGSSSNIGNNYVSWYQQLPGTAPKLLIYENNKRPSGIPDRFSGSKSGTSATLGITGLQTGDEADYYCGTWDSSLSAGGVFGTGTKVTVL |
| | CAP256 I01 | QSVLTQPPSVSAAPGQKVTISCSGSSSTIGNNYVSWYRLLPGTAPKLLIYKNDNRPSGIPDRFSGSKSGTSATLGISGLQTGDEADYYCGTWDTSLSGGGVFGTGTKVTVL |
| | CAP256 I03 | QSVLTQPPSVSAAPGQKVTISCSGSSSNIGNNFVSWYQQRPGTAPKILIYENNKRPSETPDRFSGSKSGTSATLAITGLQTADEAEYYCATWSASLSSARVFGTGTRITVL |
| | CAP256 I04 | QPVLTQPPSVSAAPGQKVTISCSGSSSNIGNNFVSWYQQRPGTAPSLLIYETNKRPSGIPDRFSGSKSATSATLAITGLQTGDEADYYCATWAASLTSARVFGTGTKVIVS |
| | CAP256 I07 | QAVLTQPPSVSAAPGQNVTISCSGSGSNIGNNFVSWYQQRPGTAPKLLIYESNKRPSGIPDRFSGSKSGTSATLAITGLQTGDEAYYYCATWAARLNSARVFGTGTMVTVL |
| | CAP256 I10 | QSVLTQPPSVSAAPGQKVTISCSGSSSNIGDNYVSWYQHLPGTAPKLLIYENTRRPSGIPDRFSGSKSGTSATLAITGLQTGDEADYYCGTWDVRPNRGAVFGTGTKVTVL |
| | CAP256 Mature | QSVLTQPPSVSAAPGQKVTISCSGNTSNIGNNFVSWYQQRPGRAPQLLIYETDKRPSGIPDRFSASKSGTSGTLAITGLQTGDEADYYCATWAASLSSARVFGTGTKVIVL |
